# Cerebellar, parietal, and frontal cortex exert distinct causal influences on cortical population codes supporting working memory

**DOI:** 10.64898/2026.05.14.724968

**Authors:** James A. Brissenden, Michael Vesia, Taraz G. Lee

## Abstract

Working memory, the transient maintenance and manipulation of information, is fundamental to human cognition. While working memory is typically thought to rely on frontoparietal cortex, the cerebellum communicates with association cortex through closed-loop cerebro-cerebellar pathways and is well positioned to influence cortical dynamics supporting cognition. It is currently unknown whether cerebellar processing is causally necessary for the maintenance of visual information and whether cortical population coding is dependent on cerebellum. We combined functional magnetic resonance imaging (fMRI) and transcranial magnetic stimulation (TMS) of frontal, parietal, and cerebellar cortex in humans. Using a generative model of population receptive fields, we show that cerebellar perturbation impairs spatial working memory recall and broadly degrades cortical spatial tuning. Direct comparisons revealed dissociable effects on population coding: cerebellar perturbation preferentially reduced stimulus-specific gain in early visual cortex, parietal perturbation preferentially suppressed stimulus-independent visual responses, and frontal perturbation preferentially increased response variability across higher-order visual, parietal, and frontal regions. These findings demonstrate that intact cerebellar function is necessary for maintaining cortical working memory representations and that cerebellar, parietal, and frontal nodes support working memory fidelity via distinct mechanisms.

**Significance Statement:** Working memory is a central process in cognition that allows us to maintain information in mind for decision-making, planning, and action. Working memory is commonly thought to rely on interactions among frontal, parietal, and sensory cortices. By combining transcranial magnetic stimulation with fMRI and model-based analyses, we show that the human cerebellum, most often associated with motor function, is also causally necessary for maintaining visuospatial information. Perturbing cerebellar, parietal, and frontal regions all impaired working memory recall, but had distinct impacts on cortical population coding underlying the maintenance of information. These findings demonstrate that prevailing theories of working memory must expand to include a distributed cortico-cerebellar network with dissociable regional contributions to the fidelity of maintained representations.

## Introduction

Working memory (WM) enables the transient maintenance of task-relevant information, supporting flexible, goal-directed behavior.^1^ Although extensive prior work has identified a distributed network spanning frontal, parietal, and visual cortices underlying visuospatial WM maintenance,^2–12^ the causal mechanisms that sustain mnemonic representations remain incompletely understood.

Traditional models posit that persistent activity in prefrontal and parietal cortices maintains visual representations over short delays.^2,3,13–15^ More recent work demonstrates that sensory cortices also contribute to WM maintenance, with stimulus-specific patterns observed during delay periods even in the absence of visual input.^7,16–19,20,21,22^ However, identifying where mnemonic information can be decoded does not reveal the circuit mechanisms that underlie the maintenance of these representations. The cerebellum is a candidate regulator of this process. Non-motor cerebellar regions are embedded in reciprocal loops with association cortex,^20^ are recruited during working memory tasks,^21,22^ and contain stimulus-specific delay-period representations.^23^ Yet prior neuroimaging evidence cannot determine whether cerebellar activity is computationally necessary for maintaining WM representations or whether it reflects activity transmitted from interconnected cortical areas.^24,25^ A causal test requires perturbing cerebellar function and then assessing how cortical population codes change as a result.

The cerebellum’s uniform microcircuitry and extensive reciprocal projections with association cortex are thought to implement forward internal models that predict the sensory consequences of actions.^26–30^ Such predictive computations may generalize beyond movement to cognition, providing a mechanism for stabilizing internal representations against noise or distraction. Optogenetic studies in rodents have shown that perturbing cerebellar output abolishes selective preparatory ramping in frontal cortex during delayed association tasks.^31–33^ These findings indicate that cerebellar output is critical for the emergence of cortical selectivity. Consistent with this account, recent computational modeling work coupled a recurrent cortical network with a feedforward, cerebellar-like module and found that feedback from this module was critical for sustaining stimulus representations across the delay period in the absence of sensory input signals.^34^ These studies raise the intriguing possibility that cerebellar computations causally support the cortical activity underlying the maintenance of working memory representations in humans.

Here we combine transcranial magnetic stimulation (TMS), fMRI, and computational modelling to test the causal contribution of the cerebellum to spatial WM in human participants. By integrating causal perturbation and model-based fMRI, this work moves beyond correlational evidence to delineate the causal computational role of cerebellar circuits in cognition. If cerebellar computations are causally necessary for the maintenance of cortical WM representations, then cerebellar perturbation should impair spatial recall, reduce the fidelity of decoded delay-period representations, and alter the underlying components of voxel spatial tuning. Comparing these effects with those produced by parietal and frontal stimulation further allowed us to determine whether different nodes support working memory representations through common or dissociable mechanisms. We find that cerebellar TMS induces WM deficits and broadly degrades cortical spatial representations. These findings identify the cerebellum as a necessary component of human working memory circuitry and show that cerebellar and frontoparietal regions make dissociable contributions to cortical population coding.

## Results

We combined TMS and fMRI to test whether the cerebellum contributes causally to the maintenance of spatial working memory. Participants completed a multi-session protocol. A baseline fMRI session collected population receptive field (pRF) mapping, resting-state scans, and anatomical MRI. The pRF data served two purposes: it allowed us to (1) define individualized TMS targets using functional connectivity and retinotopic constraints, and (2) train a spatial encoding model for subsequent analysis of working memory representations. Four additional sessions involved the application of an inhibitory TMS protocol, continuous theta-burst stimulation (cTBS), over retinotopically organized regions in cerebellar lobule VIIb (CB), intraparietal sulcus (IPS0), and superior precentral sulcus (sPCS), or a somatosensory control site just prior to performing a spatial working memory task in an fMRI scanner. During the TMS-fMRI sessions, participants performed a spatial working memory task (Fig. 1a) in which a single stimulus had to be remembered over a delay period. To improve sensitivity to TMS effects on behavior, we performed a follow-up psychophysics experiment with 4 additional sessions outside the scanner with a shortened delay (1 s vs. 8.45 s), higher set size (2 items vs. 1 item), and substantially increased trial count (480 trials vs 120 trials) (Fig. 1b), thus increasing measurement precision and power to resolve session-dependent differences.

**Figure 1.**
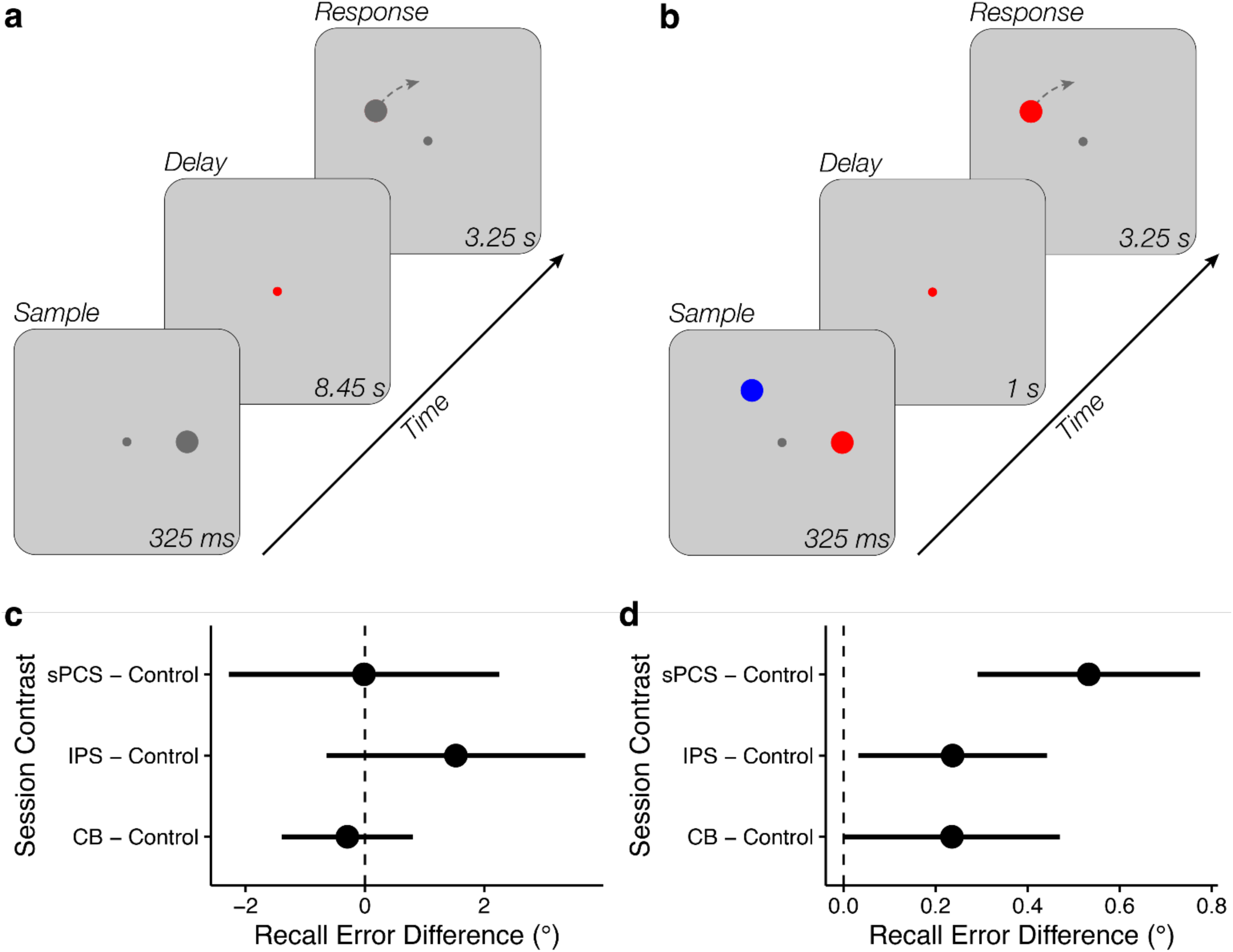
Spatial working memory (WM) paradigms and stimulation induced recall deficits. (a) Inside an fMRI scanner (Experiment 1), participants were presented with a disc stimulus (0.75°) at 4° eccentricity and were instructed to remember the location. 325 ms following the offset of the stimulus presentation period, participants were cued as to whether the trial was an ‘active’ or ‘drop’ trial (120 active trials / 20 drop trials). On ‘active’ trials, participants were instructed to maintain the location of the sample stimulus over the subsequent 8.45 s delay period, while on ‘drop’ trials they were instructed to drop the stimulus from memory and make a random response. The long delay period allowed us to isolate the BOLD response associated with WM maintenance. Following the delay period, participants were given 3.25 s to adjust a probe disc stimulus to match the remembered location. The initial angle of the probe stimulus was randomized with respect to the memorized location, which ensured that participants were unable to prospectively plan their upcoming motor response (i.e., direction of rotation) during the delay period. Thus, delay-period activity can be inferred to relate to visuospatial WM maintenance rather than preparatory motor activity. (b) In a follow-up psychophysics experiment, participants performed a similar WM task with a larger set size (2 items) and a shorter delay-period (1 s; 480 total trials). A retroactive cue presented 325 ms following the offset of the sample display indicated which of the two stimuli was to be maintained over the delay period. The probe stimulus was again randomized to prevent prospective motor planning. (c,d) Cortico-cerebellar TMS produces WM recall deficits. Parietal stimulation reliably increased recall error in the fMRI experiment (c), while cerebellar, parietal, and frontal stimulation increased recall error in the higher-powered psychophysics experiment (d).

To characterize the effects of TMS on mnemonic precision and underlying neural coding, we combined analyses of behavior with two complementary model-based fMRI approaches. Behavioral performance was evaluated in terms of spatial recall error across stimulation sites. fMRI data were modeled with a spatial encoding framework that predicted voxel-wise BOLD responses from population receptive field parameters derived from the baseline mapping session. This model was then used to (1) decode remembered locations from distributed activation patterns and the uncertainty with which those locations were represented, and (2) quantify session-dependent modulation of voxel spatial tuning, including changes in stimulus-independent responsivity, stimulus-specific gain, tuning width, and residual variance. Together, these analyses tested whether cerebellar, intraparietal sulcus, or frontal cortex perturbation altered the fidelity of spatial working memory representations across the brain and whether the stimulated sites produced dissociable effects on the underlying components of population coding.

### Cerebellar, parietal, and frontal stimulation impair mnemonic precision

We tested whether perturbing cerebellar or frontoparietal sites altered spatial working memory precision. Session effects were assessed with hierarchical within-subject permutation tests and FDR correction.

In the TMS-fMRI experiment (Experiment 1; 120 active trials/session), IPS stimulation increased recall error relative to control (IPS – Control = 1.54°, *p* = 0.005 corrected). Spatial recall was largely unchanged for CB and sPCS stimulation (CB – Control = –0.304°, *p* = 0.76 corrected; sPCS – Control = –0.025°, *p* = 0.96 corrected). The deleterious effect of IPS stimulation on behavior remained significant after regressing out session order (*p* = 0.005 corrected).

Although the single-item WM task we employed in the scanner was ideal for fMRI analyses, the low trial count (120 total) and low memory load may have impaired our ability to detect behavioral deficits following TMS. In the TMS-psychophysics experiment with higher statistical power (Experiment 2; 480 active trials/session), cerebellar, parietal and frontal stimulation all resulted in increased recall error relative to control stimulation (sPCS – Control = 0.525°, p = 0.0006 corrected; CB – Control = 0.221°, p = 0.008 corrected; IPS – Control = 0.225°, p = 0.008 corrected). The cerebellar and sPCS effects remained significant after controlling for session order (sPCS – Control: *p* = 0.0006 corrected; CB – Control: *p* = 0.04 corrected; IPS – Control: *p* = 0.13 corrected).

Across the two experiments, we observe behavioral deficits associated with each targeted region: IPS in Experiment 1, and cerebellum and sPCS in Experiment 2, where the larger trial count provided greater sensitivity to detect changes in spatial recall precision.

### Cerebellar and frontoparietal stimulation reduce the fidelity of spatial working memory representations

To assess how perturbing different nodes of the cortico-cerebellar network influenced the fidelity of spatial working memory representations, we applied a generative probabilistic decoding approach.^23,38–40^ This analysis uses voxelwise spatial tuning parameters (Fig. 2) derived from the baseline pRF session to estimate, for each trial, the posterior probability of each possible remembered location (0°–359° at 4° eccentricity) given the observed multivoxel activation pattern during the memory delay (Fig. 3). Averaging trial-aligned posterior distributions produces a session-level probability density over stimulus locations for each region of interest (ROI). Sharper peaks around 0° reflect better decoding of the remembered spatial location. Control stimulation consistently produced more sharply peaked distributions relative to CB, IPS, and sPCS stimulation (see Fig. 4a). From trial-level posterior distributions, we derive two measures: estimation efficiency, the inverse of mean circular squared error between the posterior mean and the true stimulus, and decoded uncertainty, the mean posterior standard deviation across trials.

**Figure 2.**
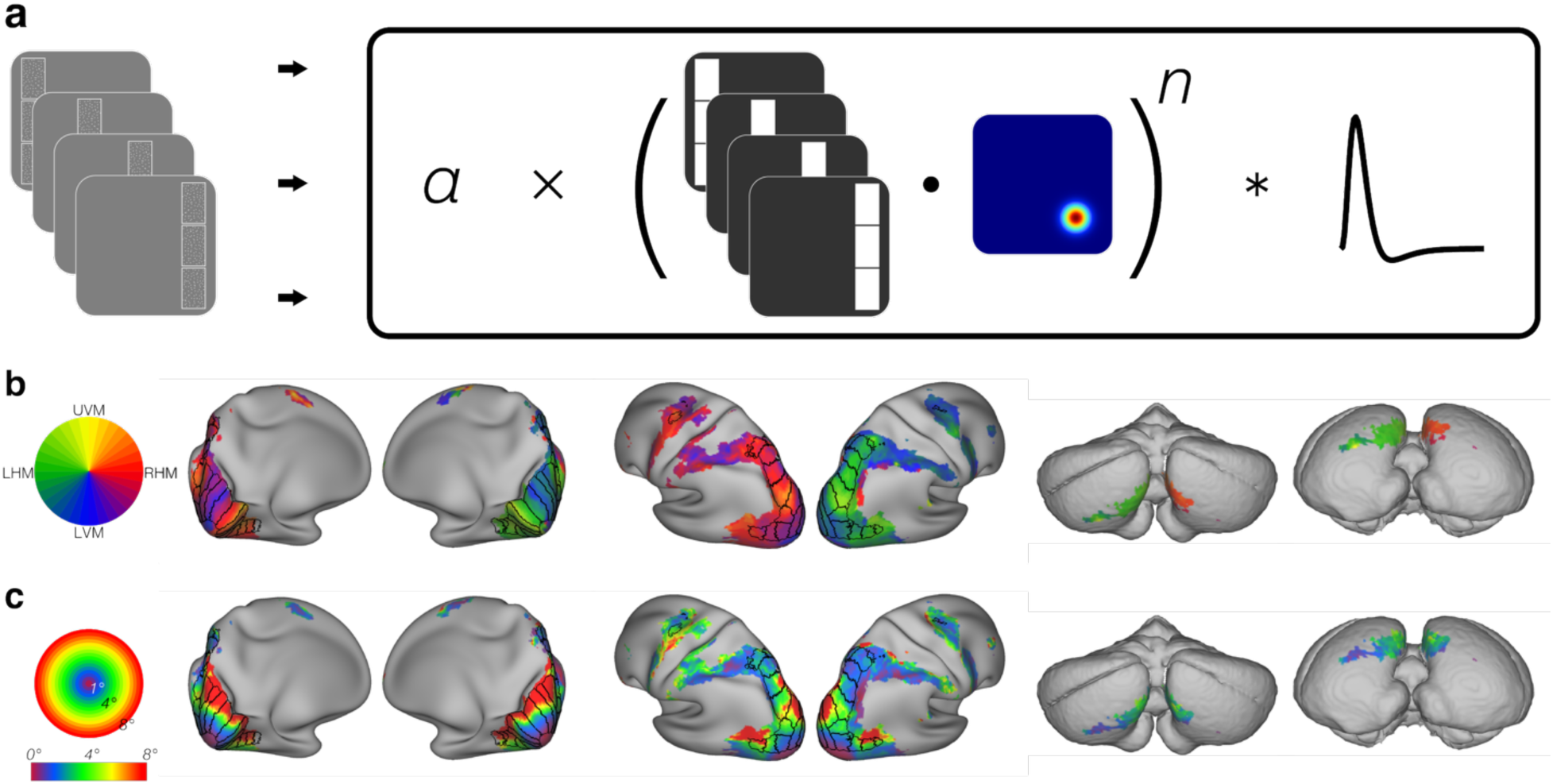
Population receptive fielding (pRF) mapping. (a) In an initial fMRI session, participants performed an attentionally demanding visual field mapping task which required discriminating and comparing the direction of motion of three equal sized patches of moving dot patterns at different positions within the display.^35^ Resulting BOLD responses were modeled with compressive spatial summation model.^36^ This model produces a 2D Gaussian receptive field for each voxel (x, y, σ) along with gain (a) and compressive nonlinearity (n) parameters. (b) Group average (N = 27) polar angle preferences across cortex and cerebellum. These results replicate prior work demonstrating spatial selectivity in cerebellum.^22,37^ (c) Group average eccentricity preferences across cortex and cerebellum.

**Figure 3.**
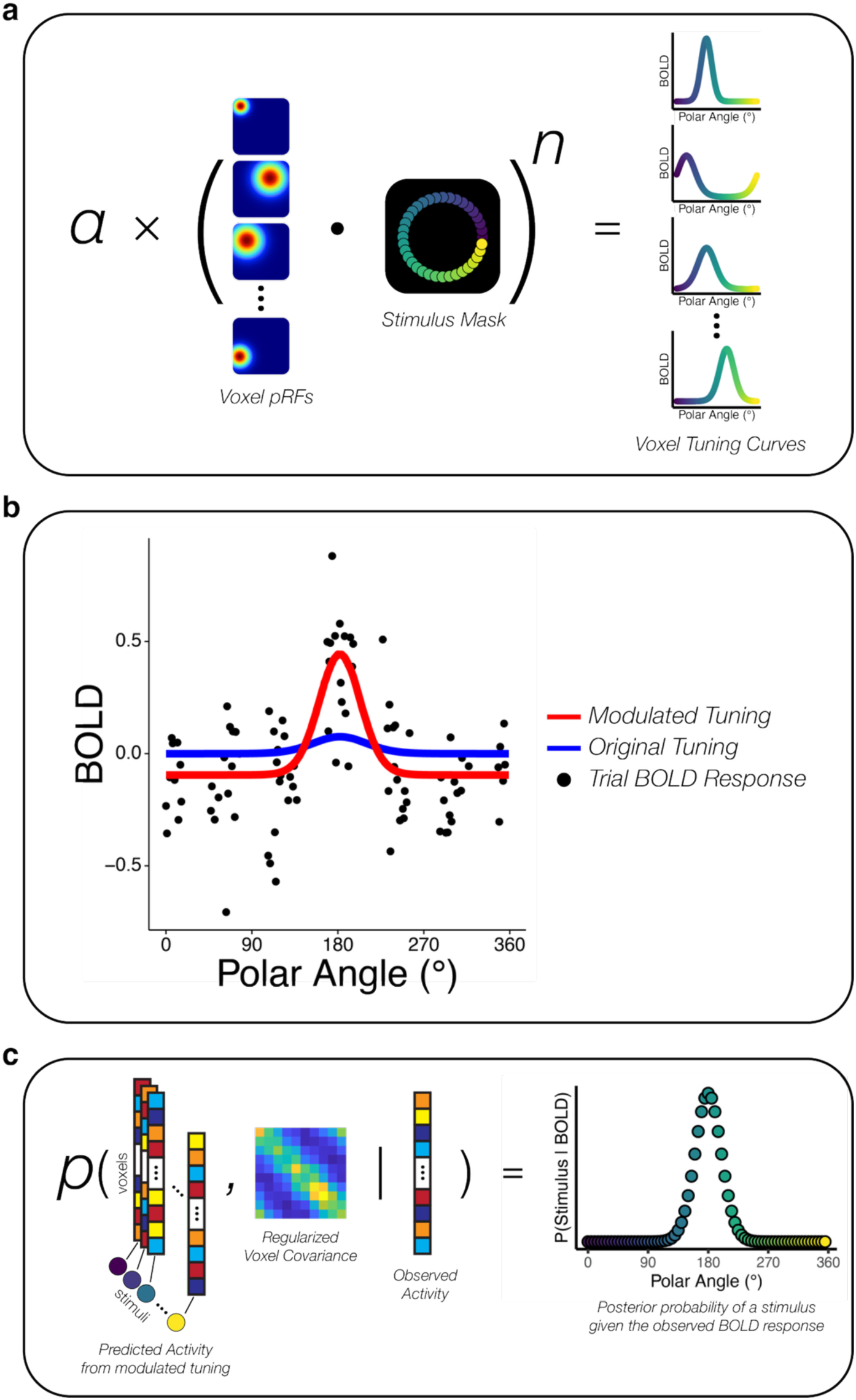
Probabilistic spatial encoding model. (a) fMRI data were modeled with a spatial encoding framework that predicted voxelwise BOLD responses from population receptive field parameters derived from the baseline mapping session (Fig. 2a) and stimulus masks corresponding with all possible spatial WM locations probed in the TMS-fMRI sessions. (b) Spatial tuning curve modulation for an example voxel. The model accounted for session-specific modulation of spatial tuning curves. Each black dot represents the late-delay BOLD response for a single WM trial. The blue curve represents the predicted response to each spatial location from the original model parameters, while the red curve reflects an adjustment in gain, width, and baseline activation as a function of task. (c) Probabilistic decoding of spatial location. For each trial, we computed the posterior probability of each possible stimulus location given the observed multi-voxel BOLD activity on that trial, while taking into account covariance across voxels (see methods).

**Figure 4.**
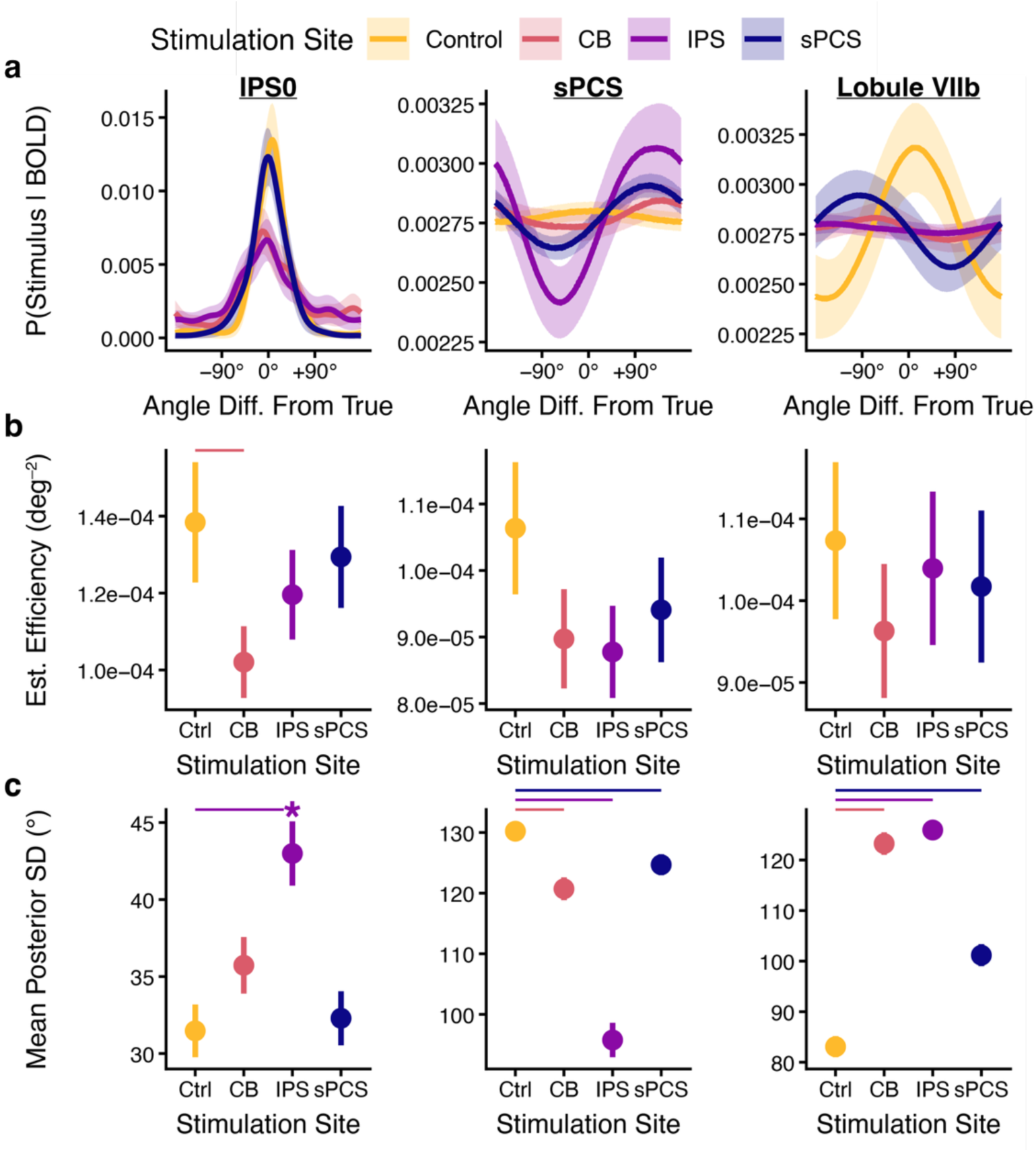
Cortico-cerebellar stimulation degrades spatial decoding across multiple brain regions. (a) Mean posterior probability distribution. Averaging trial-aligned posterior distributions produces a session-level probability density over stimulus locations for each region of interest (ROI). Sharper peaks around 0° reflect better decoding of the remembered spatial location. (b) Estimation efficiency (inverse of mean circular squared error between the posterior mean and the true stimulus) reflecting decoding accuracy for each session and region. (c) Mean posterior standard deviation reflecting decoding uncertainty. Error bars reflect a bootstrap estimate of standard error. Horizontal bars indicate corrected p < 0.05 from a hierarchical permutation test. If at least one WM active stimulation condition differed significantly from control within a given ROI, pairwise comparisons among the WM active sites (CB versus IPS, CB versus sPCS, and IPS versus sPCS) were then performed for that ROI. Asterisks indicate a selective deficit relative to the other two WM active sites (FDR corrected p < 0.05). Each column reflects results for one ROI. IPS – intraparietal sulcus; sPCS – superior pre-central sulcus.

Cerebellar stimulation produced robust and spatially distributed reductions in the fidelity of spatial representations. Relative to control stimulation, CB stimulation significantly broadened posterior distributions in cerebellar lobule VIIb and a higher-order visual area (V3AB) known to exhibit robust brain-behavior correlations during WM^40^ (all p ≤ 0.0004 corrected; Fig. 4c). This result indicates that cerebellar perturbation increased uncertainty in the representation of remembered locations across both cortical and cerebellar regions. Similar increases in uncertainty were also observed in higher-order parietal regions IPS1 and IPS3 (Supplementary Fig. 2). In addition to these increases in decoded uncertainty, cerebellar stimulation also reduced estimation efficiency in V3AB (p = 0.02 corrected) and IPS0 (p = 0.02 corrected) (Fig. 4b, Supplementary Fig. 1b), reflecting a measurable decline in the accuracy with which remembered locations could be reconstructed from population activity. These results indicate that cerebellar perturbation broadly degrades the fidelity of spatial representations and additionally reduces reconstruction accuracy in key visual and parietal regions.

Parietal stimulation produced a comparable pattern of representational disruption, primarily expressed as increased uncertainty. IPS stimulation significantly increased decoded uncertainty in V3AB, IPS0, and cerebellar lobule VIIb (all p ≤ 0.0004 corrected; Fig. 4c; Supplementary Fig. 1c), as well as in IPS1 and IPS3 (Supplementary Fig. 2c). IPS stimulation did not significantly reduce estimation efficiency relative to control in any ROI, indicating that its principal effect on decoded representations was an increase in posterior uncertainty rather than a reduction in reconstruction accuracy.

Frontal stimulation also altered decoded representations across cortical and cerebellar regions. Relative to control, sPCS stimulation significantly increased decoded uncertainty in V3AB and cerebellar lobule VIIb (both p ≤ 0.0004 corrected; Fig. 4c; Supplementary Fig. 1c), as well as in IPS1 and IPS3 (Supplementary Fig. 2c). In contrast, no significant increase in uncertainty was observed in IPS0 (Fig. 4c). sPCS stimulation additionally reduced estimation efficiency in V3 (p = 0.01 corrected; Supplementary Fig. 1b), but not in the other ROIs examined.

The sPCS region itself exhibited an unusual decoding profile across all stimulation conditions that warrants separate consideration. Under control stimulation, sPCS exhibited weak spatial selectivity, as evidenced by high decoded uncertainty, approaching the values expected under a near-uniform posterior. Following stimulation of all sites (CB, IPS, or sPCS), decoded uncertainty in sPCS decreased relative to control. However, these narrower posterior distributions were not centered on the veridical stimulus location (Fig. 4a), resulting in numerically *worse* estimation efficiency. Thus, reduced uncertainty in sPCS did not reflect enhanced spatial selectivity but rather a qualitative change in the selectivity profile across stimulus locations.

Follow-up comparisons among the active stimulation sites revealed several regional differences in their effects on spatial decoding (Supplmentary Table 2). Estimation efficiency in V3 was lower following sPCS stimulation than following either CB or IPS stimulation (both p < 0.001 corrected), whereas estimation efficiency in IPS0 was lower following CB than sPCS stimulation (p = 0.043 corrected). Differences in decoded uncertainty were similarly region dependent. IPS stimulation produced greater uncertainty than CB and sPCS stimulation in IPS0 (p = 0.019 and p = 0.001 corrected, respectively), whereas CB and sPCS stimulation produced greater uncertainty than IPS stimulation in IPS1 (both p < 0.001 corrected). In IPS3, uncertainty was greater following sPCS than CB stimulation (p = 0.021 corrected), while in cerebellar lobule VIIb, CB and IPS stimulation produced greater uncertainty than sPCS stimulation (both p < 0.001 corrected). No significant differences among active stimulation sites were observed in V3AB.

Taken together, the decoding results indicate that perturbation of cerebellar, parietal, and frontal nodes each alters the fidelity of spatial working memory representations across distributed cortical and cerebellar regions. These effects were expressed predominantly as changes in decoded uncertainty, with more spatially restricted effects on estimation efficiency. Moreover, direct comparisons among the active stimulation sites revealed region-specific differences in both measures, indicating that the three perturbations produce distinct profiles of representational change rather than differing simply in overall effect magnitude. These findings motivate further examination of how TMS alters the underlying spatial tuning properties of voxel responses.

### Cortico-cerebellar TMS alters voxel-level tuning across the cortico-cerebellar network

Probabilistic decoding revealed how stimulation altered the fidelity of remembered spatial representations, but it does not identify which properties of the underlying voxel response profiles were altered by perturbation. Reduced decoding performance could arise from weaker stimulus-specific responses, broader spatial tuning, greater response variability, or some combination of these factors. Moreover, decoding does not capture stimulus-independent changes in responsivity, which may nevertheless reveal important effects of stimulation on cortical population responses. We therefore examined how cerebellar, parietal, and frontal stimulation modulated voxel-level response parameters to determine whether perturbation of these regions produced common or dissociable signatures during working memory maintenance. This analysis separately quantified multiplicative gain (*ϕ*_*gain*_), tuning width (*ϕ_size_*), additive baseline shifts (*ϕ_baseline_*), and residual variance (*ϕ_var_*) relative to the baseline pRF-predicted response (Fig. 5). For gain, width, and variance, values greater than 1 indicate increased estimates relative to the tuning estimated in the baseline session, whereas values less than 1 reflect reductions. For the baseline parameter, positive values indicate increases relative to the baseline and negative values reflect decreases. However, as the compressive spatial summation pRF model produces only positive response values, note that negative baseline modulation values do not suggest a true reduction from the baseline session. Rather, they reflect the model’s adjustment to z-scored time series with zero mean, as the BOLD response during working memory maintenance has a smaller magnitude than the response to visual stimulation and motor responses.

**Figure 5.**
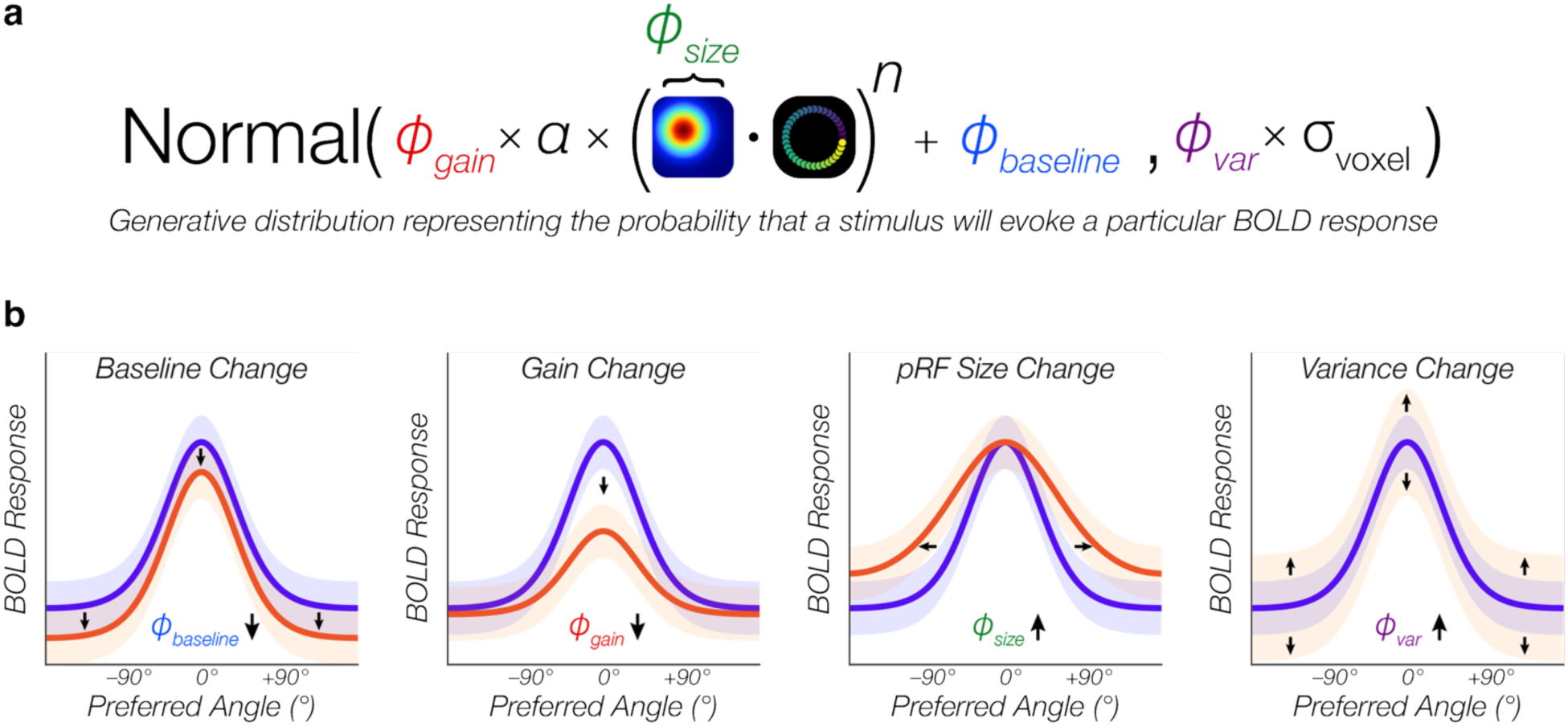
Spatial tuning curve modulation and hypothetical effects of stimulation. (a) Analysis separately quantified TMS-induced changes in multiplicative gain (ϕ_gain_), tuning width (ϕ_size_), additive baseline shifts (ϕ_baseline_), and residual variance (ϕ_var_) relative to the baseline pRF-predicted response. Gaussian center (x, y) and compressive nonlinearity exponent (n) were held constant. (b) Possible effects of stimulation on different aspects of voxel spatial tuning.

Cerebellar stimulation produced widespread changes across multiple components of voxel spatial tuning. Relative to control stimulation, cerebellar TMS significantly reduced the additive baseline parameter in all ROIs (all p ≤ 0.003 corrected; Figs. 6-8; Supplementary Table 3), indicating a broad reduction in stimulus-independent BOLD activity. Because the baseline term captures components of the response that do not vary with the spatial location of the stimulus, these shifts reflect a global suppression of visuocortical activity consistent with prior demonstrations that cTBS reduces neural responsiveness across cortical^41–43^ and cerebellar targets.^44–48^ In addition to these baseline effects, cerebellar stimulation reduced stimulus-specific gain in all visual, parietal, and cerebellar regions (all p ≤ 0.003 corrected), increased tuning width in IPS3 and VIIb (p ≤ 0.03 corrected), and increased residual variance in V2, IPS0-1, IPS3, sPCS, and VIIb (all p ≤ 0.01 corrected). (Figs. 6-8; Supplementary Table 3).

**Figure 6.**
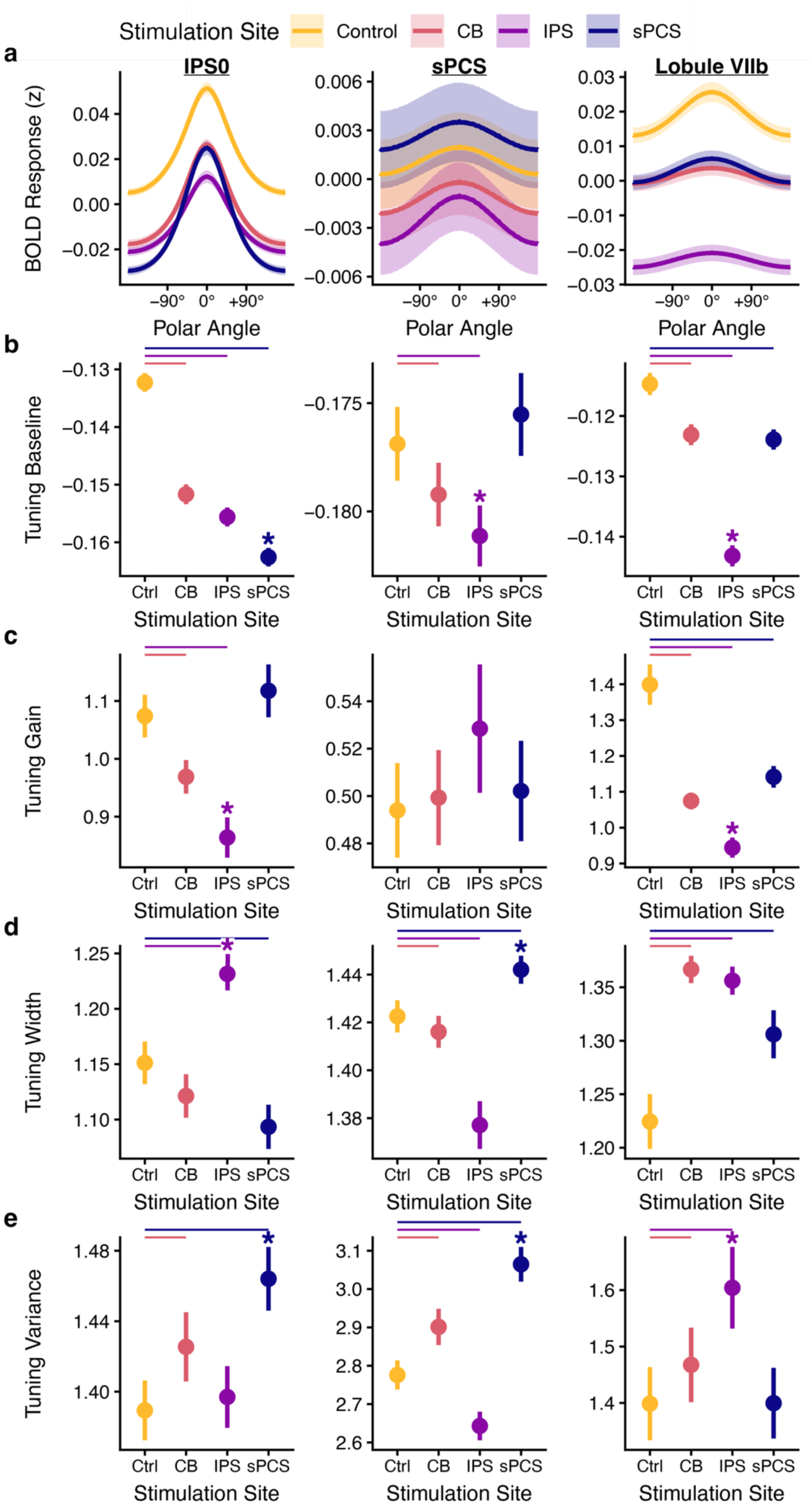
TMS induced changes in spatial tuning during WM maintenance. (a) Mean tuning curve across voxels for each stimulation session within each ROI. For visualization, the tuning curves are demeaned across sessions. Shaded ribbons reflect a bootstrap estimate of SEM across voxels. (b) Spatial tuning curve baseline for each session and ROI. This value reflects stimulus-independent BOLD activity during the delay-period. (c) Spatial tuning curve gain for each session and ROI. This value reflects the BOLD response for preferred stimulus locations relative to non-preferred locations. (d) Spatial tuning curve width for each session and ROI. (e) Spatial tuning curve variance for each session and ROI reflecting the distribution of BOLD responses for repeated presentations of the same stimulus. Error bars across b-e reflect a bootstrap estimate of SEM. Horizontal bars indicate corrected p < 0.05 from a hierarchical permutation test. If at least one WM active stimulation condition differed significantly from control within a given ROI, pairwise comparisons among the WM active sites (CB versus IPS, CB versus sPCS, and IPS versus sPCS) were then performed for that ROI. Asterisks indicate a selective deficit relative to the other two WM active sites (FDR corrected p < 0.05).

**Figure 7.**
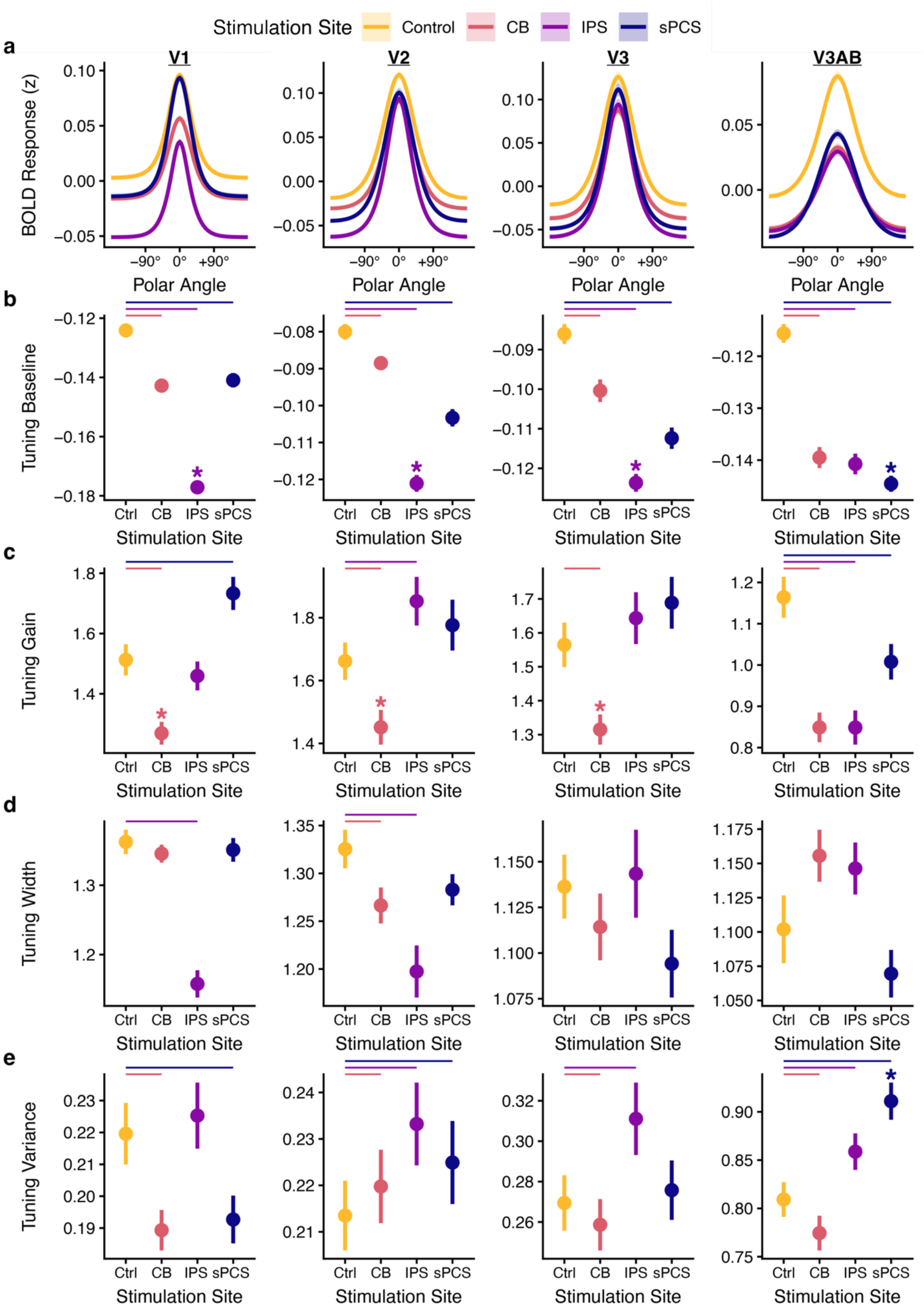
TMS induced changes in visual cortex spatial tuning during WM maintenance. (a) Mean tuning curve across voxels for each stimulation session within each ROI. For visualization, the tuning curves are demeaned across sessions. Shaded ribbons reflect a bootstrap estimate of SEM across voxels. (b) Spatial tuning curve baseline for each session and ROI. This value reflects stimulus-independent BOLD activity during the delay-period. (c) Spatial tuning curve gain for each session and ROI. This value reflects the BOLD response for preferred stimulus locations relative to non-preferred locations. (d) Spatial tuning curve width for each session and ROI. (e) Spatial tuning curve variance for each session and ROI reflecting the distribution of BOLD responses for repeated presentations of the same stimulus. Error bars across b-e reflect a bootstrap estimate of SEM. Horizontal bars indicate corrected p < 0.05 from a hierarchical permutation test. If at least one WM active stimulation condition differed significantly from control within a given ROI, pairwise comparisons among the WM active sites (CB versus IPS, CB versus sPCS, and IPS versus sPCS) were then performed for that ROI. Asterisks indicate a selective deficit relative to the other two WM active sites (FDR corrected p < 0.05).

**Figure 8.**
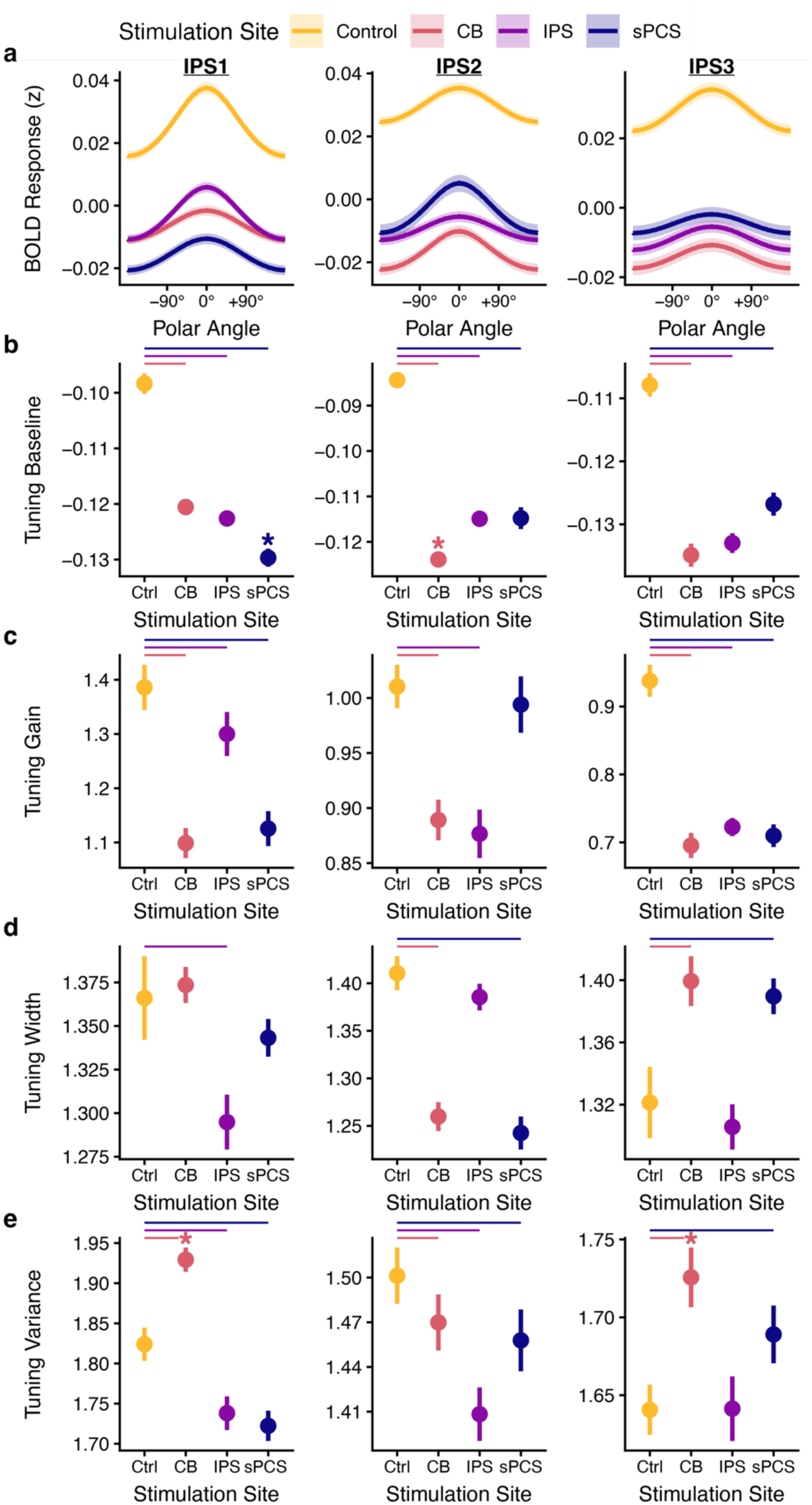
TMS induced changes in parietal cortex spatial tuning during WM maintenance. (a) Mean tuning curve across voxels for each stimulation session within each ROI. For visualization, the tuning curves are demeaned across sessions. Shaded ribbons reflect a bootstrap estimate of SEM across voxels. (b) Spatial tuning curve baseline for each session and ROI. This value reflects stimulus-independent BOLD activity during the delay-period. (c) Spatial tuning curve gain for each session and ROI. This value reflects the BOLD response for preferred stimulus locations relative to non-preferred locations. (d) Spatial tuning curve width for each session and ROI. (e) Spatial tuning curve variance for each session and ROI reflecting the distribution of BOLD responses for repeated presentations of the same stimulus. Error bars across b-e reflect a bootstrap estimate of SEM. Horizontal bars indicate corrected p < 0.05 from a hierarchical permutation test. If at least one WM active stimulation condition differed significantly from control within a given ROI, pairwise comparisons among the WM active sites (CB versus IPS, CB versus sPCS, and IPS versus sPCS) were then performed for that ROI. Asterisks indicate a selective deficit relative to the other two WM active sites (FDR corrected p < 0.05).

Parietal stimulation also produced widespread tuning changes. IPS stimulation produced robust reductions in baseline activity across all ROIs (all p = 0.0002 corrected; Figs. 6-8; Supplementary Table 3), consistent with a global suppression of stimulus-independent responses. Beyond these stimulus-independent baseline effects, IPS stimulation also significantly reduced stimulus-dependent gain in V3AB, parietal cortex (IPS0-3), and lobule VIIb (all p ≤ 0.04 corrected), increased tuning width in IPS0 and lobule VIIb (both p = 0.008 corrected), and increased residual variance in extrastriate visual cortex (V2-3, V3AB) and lobule VIIb (all p = 0.0003 corrected) (See Supplementary Table 3).

Frontal stimulation significantly reduced baseline activity in all regions except itself (all p = 0.0002 corrected), and reduced stimulus-specific gain in V3AB, IPS1, IPS3, and cerebellar lobule VIIb (all p ≤ 0.003 corrected). Tuning width was increased in IPS3, sPCS, and lobule VIIb (p ≤ 0.03 corrected), and residual variance increased in V2, V3AB, IPS0, IPS3, and sPCS itself (all p ≤ 0.002 corrected).

Although stimulation to cerebellar, parietal, and frontal cortex all produced robust and widespread effects on spatial tuning throughout the brain, there were three prominent dissociations in their effects on voxel tuning during WM maintenance (Figs. 6-8; Supplementary Table 4). First, cerebellar stimulation produced greater reductions in stimulus-specific gain than either IPS or sPCS stimulation throughout early visual cortex (*ϕ*_*gain*_; V1–V3; all p < 0.001 corrected; Fig. 7). Second, IPS stimulation produced larger decreases in stimulus-independent BOLD activity than either cerebellar or sPCS stimulation across these same visual regions (*ϕ_baseline_*, V1–V3; all p < 0.001 corrected; Fig. 7). Third, sPCS stimulation produced greater increases in residual response variance than either cerebellar or IPS stimulation in V3AB, IPS0, and sPCS itself (*ϕ_var_*; all p ≤ 0.013 corrected; Figs. 6 and 7). Thus, although perturbation of each WM brain area altered multiple tuning parameters, TMS to each area exhibited dissociable signatures during the WM delay period: cerebellar perturbation preferentially suppressed stimulus-specific visual cortex gain, parietal perturbation preferentially reduced the stimulus-independent component of visual cortex responses, and frontal perturbation preferentially increased response variability across higher-order visual, parietal, and frontal regions. These site-specific signatures indicate that cerebellar, parietal, and frontal nodes support WM maintenance through partly distinct influences on the strength, overall responsivity, and reliability of cortical population responses.

## Discussion

We provide evidence that the human cerebellum, along with frontoparietal cortex, contributes causally to the maintenance of spatial working memory representations. We combined individualized cTBS with functional neuroimaging and model-based analyses to determine how perturbing cerebellar, parietal, and frontal nodes affects behavior and population coding across a distributed cortico-cerebellar working memory network. Across two experiments, stimulation of each working memory active site impaired spatial recall. Probabilistic decoding further showed that stimulation reshaped delay-period representations of remembered location across cortical and cerebellar regions. Decomposing these effects into voxel-level tuning components revealed dissociable signatures across stimulation sites: cerebellar perturbation preferentially suppressed stimulus-specific gain in early visual cortex, parietal perturbation preferentially reduced the stimulus-independent component of visual cortical responses, and frontal perturbation preferentially increased response variability across higher-order visual, parietal, and frontal regions. These findings establish the cerebellum as a necessary component of human working memory circuitry and show that cerebellar, parietal, and frontal regions can support the fidelity of cortical population codes through distinct mechanisms.

By demonstrating that perturbation of frontal and parietal cortices reduces the fidelity of spatial working memory representations, our results are consistent with a large body of work implicating frontoparietal regions in working memory using both correlational and causal approaches.^2,3,5,6,10–13,49,50^ Prior work has also implicated the cerebellum in working memory, but the evidence has been predominantly correlational. Neuroimaging and neuropsychological studies have associated cerebellar activity with verbal working memory and articulatory rehearsal processes, particularly in lobules VI and Crus I/II,^51–54^ while a separate body of work has linked the posterior cerebellum to visual and spatial working memory demands.^21–23,55–58^ Convergent large-scale functional mapping has further indicated that cerebellar areas are systematically coupled with association networks supporting cognitive control, attention, and memory.^59–64^ Yet despite this consistency across studies, such findings cannot determine whether cerebellar activity is necessary for working memory maintenance or instead reflects the downstream consequences of cortical processing. By combining cerebellar stimulation with model-based fMRI measures of representational fidelity, the present study provides causal evidence that cerebellar output contributes to the maintenance of spatial working memory representations in humans.

In the current study, behavioral performance was reliably affected by theta-burst stimulation. TMS disruption of cerebellar, parietal, and frontal targets each increased recall error relative to control. These results establish that perturbing any node in this cortico-cerebellar circuit can impair spatial recall. Probabilistic decoding revealed corresponding changes in the neural representations supporting performance. Across stimulation conditions, posterior distributions over remembered location were systematically reshaped, indicating changes in the reliability with which delay-period population activity specified spatial position. Direct comparisons among the active stimulation sites revealed regional differences in these decoding effects, but did not point to a clear dissociable pattern of influence on working memory encoding across the brain.

However, a TMS-induced reduction in decoding performance does not distinguish whether this deficit reflects weaker stimulus-specific responses, more diffuse stimulus preferences, greater response variability, or some combination of these factors. Moreover, because spatial decoding depends on stimulus-dependent differences in population responses, it is insensitive to additive changes in stimulus-independent responsivity that may nevertheless reflect robust causal effects of stimulation. We therefore complemented the decoding analysis with a voxel-level tuning modulation analysis that separated changes in stimulus-specific gain, tuning width, response variability, and stimulus-independent responsivity. The disproportionate reduction of visual cortical gain following cerebellar stimulation suggests that cerebellar output helps sustain the stimulus-specific modulation of sensory populations during memory maintenance. By contrast, the stronger reduction of the stimulus-independent visual cortex response following parietal stimulation is consistent with the idea that parietal cortex exerts a broad influence on overall task responsivity across cortex. Frontal stimulation most strongly increased residual response variance, suggesting a contribution to the reliability of population responses across repeated representations of the same stimulus. Because all three stimulation sites affected multiple parameters, these signatures should not be interpreted as exclusive functions. Nevertheless, direct comparisons demonstrate that cerebellar, parietal, and frontal nodes exert distinguishable causal influences on cortical population codes during working memory maintenance.

The preferential reduction in visual cortical gain following cerebellar stimulation provides a more specific account of how cerebellar output may support distributed working memory representations. Recent causal evidence from animal models provides a useful framework for interpreting this result. Perturbing cerebellar output can directly reshape cortical dynamics and task-related activity patterns, indicating that cerebellar circuits actively influence forebrain computations rather than merely refining movement execution.^31–33,65^ For example, disrupting cerebellar nuclei abolishes the emergence of selective preparatory dynamics in frontal cortex during a delayed association task,^32^ demonstrating that cerebellar output can be necessary for the emergence and maintenance of task-relevant cortical states. More recent work in macaques similarly found that blocking cerebellar outflow altered preparatory cortical activity, increasing its dimensionality and reducing generalization across movement conditions.^66^ Although these studies did not quantify multiplicative gain in the formal sense used here, they converge on the idea that cerebellar output helps preserve the selectivity and structure of task-relevant cortical activity. Yet, due to the fact that many of these paradigms tightly couple working memory demands with prospective motor planning, it is not possible to ascertain whether cerebellar computations support the maintenance of non-motor representations. Here, by randomizing the initial location of the probe stimulus, we eliminate the opportunity for subjects to pre-plan a specific sequence of motor actions during the delay. Under these conditions, cerebellar stimulation reduced both the fidelity of delay-period spatial representations and stimulus-specific gain in visual cortex, indicating that cerebellar influences on cortical selectivity extend beyond motor planning or response selection.

How might cerebellar computations support working memory? Prominent theories propose that the cerebellum implements forward internal models that help predict and stabilize neural dynamics.^27–30,34^ Consistent with this role, the cerebellum has been extensively implicated in visuomotor adaptation, where it is thought to update internal models based on sensory prediction errors.^26,27,29,67–72^ In a working memory context, this view implies that cerebellar output could contribute to maintaining task-relevant cortical states by reducing drift, limiting noise accumulation, or coordinating distributed population codes across frontoparietal and sensory regions. Computational work is consistent with this idea: coupling a recurrent cortical network to a cerebellar-like module improves the stability of stimulus representations across delays without sensory input.^34^ These results support the view that cerebellar computations help sustain high-fidelity cortical representations during memory maintenance. Together with the present reduction in stimulus-specific visual gain, these findings support the view that cerebellar computations help sustain the selectivity and stability of cortical representations during memory maintenance.

The dissociable effects of parietal and frontal stimulation are also consistent with prior evidence that these regions exert distinct influences on visual cortex. In concurrent TMS-fMRI studies, stimulation of a frontal site identified as the human frontal eye field (FEF), near the superior precentral sulcus, produced an eccentricity-dependent modulation of retinotopic visual cortex, increasing BOLD responses for peripheral visual-field representations while decreasing responses for central representations in V1–V4. These effects were observed regardless of whether visual input was present or absent, leading the authors to conclude that frontal cortex influences visual cortex in a top-down manner that is largely independent of concurrent bottom-up drive.^73^ In contrast, IPS stimulation increased activity across V1–V4 primarily in the absence of visual stimuli, with effects that were relatively uniform across eccentricity. The dependence of these effects on visual context suggests that parietal feedback to early visual cortex may be most apparent when bottom-up visual drive is weak or absent, whereas strong sensory input may reduce the influence of parietal feedback on these regions.^74^ Although a one-to-one correspondence with the present findings should not be expected given substantial differences in stimulation protocols and behavioral tasks, these studies provide causal evidence that parietal and frontal regions can modulate visual cortex through distinct patterns of influence. A related functional dissociation has also been observed during a working memory guided saccade paradigm. Mackey and Curtis applied TMS to topographically organized sPCS and IPS2 during the memory delay and found differential patterns of behavioral impairment. sPCS stimulation selectively increased errors in the initial memory-guided saccade while leaving the final corrected eye position unaffected, whereas IPS2 stimulation primarily impaired the final eye position. They interpreted this asymmetric pattern as evidence that sPCS preferentially supports prospective action-related representations, whereas IPS contributes more strongly to retrospective representations of the remembered sensory location.^75^ Our results extend these findings by identifying distinct components of population responses affected by each site. The preferential reduction of stimulus-independent visual cortex responses we observed following IPS stimulation is consistent with a broad parietal influence on the overall responsivity of sensory populations. In contrast, the disproportionate increase in residual variance following sPCS stimulation suggests a contribution of frontal cortex to the reliability of distributed cortical responses. Chang et al. further showed that mean firing rate and trial-to-trial response variability in macaque FEF exhibited markedly different dynamics during a visuospatial working memory task. Firing rates remained spatially selective throughout the memory delay, whereas the Fano factor decreased broadly following visual stimulation but returned toward baseline during the delay and showed little spatial or attentional modulation. Thus, response variability did not simply track the magnitude or selectivity of neuronal responses, leading the authors to suggest that it may instead reflect shifts in the relative contributions of feedforward and recurrent excitatory drive.^76^ Although residual BOLD variance is not directly equivalent to spike-count variability, the disproportionate increase in residual variance following sPCS stimulation raises the possibility that frontal perturbation alters the recurrent or top-down inputs that normally constrain the reliability of distributed population responses during working memory maintenance.

We recently provided behavioral evidence that cerebellar adaptive control mechanisms traditionally associated with motor learning extend to spatial cognition. In a task that repeatedly induced covert attentional allocation errors, we found that spatial working memory representations gradually shifted to counteract those errors, exhibiting exponential time courses, spatially graded transfer, and rapid de-adaptation.^77^ These signatures closely parallel canonical sensorimotor adaptation. The present finding that cerebellar perturbation preferentially weakens stimulus-specific visual gain identifies a potential neural substrate through which such calibration could influence mnemonic representations. Cerebellar feedback may support not only the gradual updating of spatial representations following errors, but also their online stabilization during memory delays. Future work will be needed to determine whether these adaptive and maintenance functions arise from a common cerebellar computation.

These results inform ongoing debates regarding the neural substrates of working memory. Sensory recruitment accounts propose that mnemonic content is maintained within stimulus-selective sensory populations,^1,4,7,16,78^ whereas alternative views emphasize higher-order or distributed storage mechanisms.^9,79,80^ The present data support a distributed perspective: perturbing cerebellar, parietal, and frontal nodes altered representational fidelity, while direct comparisons revealed dissociable effects on the underlying components of cortical tuning. Together, these findings challenge cortico-centric accounts of working memory by demonstrating that the persistent maintenance of cognitive representations depends critically on cerebellar contributions to a broader maintenance network and that different nodes within this network support representational fidelity through distinguishable mechanisms.

## Methods

### Design overview

We used a within-subject, multi-session protocol to test how cerebellar, parietal, and frontal visuospatial maps support spatial working memory (WM). A baseline MRI session provided high-resolution anatomical images, resting-state fMRI, and retinotopic pRF mapping. These data served two roles: (1) defining individualized TMS targets using retinotopy and functional-connectivity constraints within atlas-based anatomical search spaces (cerebellar lobule VIIb (CB), intraparietal sulcus area IPS0 (IPS), superior precentral sulcus (sPCS), and a somatosensory control site); and (2) training a spatial encoding model for subsequent spatial decoding analyses of the spatial WM task.

Each participant then completed four TMS-fMRI sessions, one per stimulation site (CB, IPS, sPCS, control). At the start of each session, continuous theta-burst stimulation (cTBS) was delivered at an individually titrated intensity. Immediately afterward, participants performed the spatial WM task during fMRI while cTBS after-effects were present.^41^ Session order was counterbalanced across participants. The primary fMRI outcomes were decoding-based estimates of remembered location and uncertainty, and voxel-wise tuning-curve modulation. Each of these measures derived from the fMRI data following stimulating a site of interest was contrasted with that following stimulation of the control site.

A subset of participants completed additional TMS-psychophysics sessions outside the scanner (same four stimulation sites). Experiment 2 used a slightly modified WM task relative to the fMRI sessions in Experiment 1 and was more strongly powered to detect subtle behavioral effects (detailed below).

### Participants

28 total healthy adult volunteers participated in this study. All research protocols were approved by the Health Sciences and Behavioral Sciences Institutional Review Board at the University of Michigan. All participants gave written informed consent. Participants were paid $15/hr for their participation. A $50 bonus payment was disbursed if a participant successfully completed all sessions. Participants were recruited from the University of Michigan and the surrounding community. All participants possessed normal or corrected-to-normal vision.

Due to the longitudinal nature of this within-subject design and the logistics of scheduling up to 9 total sessions, many participants did not complete all possible sessions. 9 participants did not return for the TMS-fMRI sessions, but their baseline sessions were important for training the pRF model to assess the impact of TMS on spatial representation (see below). Of the combined TMS-fMRI sessions, 17 subjects completed the Control session, 17 subjects completed the sPCS session, 16 subjects completed the CB session, and 16 completed the IPS session. 14 of the original participants were invited to complete an additional 4 sessions combining TMS with psychophysics. 12 subjects completed all sessions. Of the two remaining subjects, one completed only the control session and the other completed all sessions except for the CB session.

### MRI Acquisition

MR data were acquired with a GE Signa MR750 3.0T MRI scanner with a 32-channel head coil. The baseline session collected high-resolution 3D inversion-recovery T1-weighted MPRAGE (sagittal; TR = 2523 ms; TE = 3.55 ms; TI = 1060 ms; flip angle = 8°; 0.8 mm isotropic voxels; matrix = 320 × 320; 208 slices; parallel imaging: auto-calibrating reconstruction for cartesian imaging (ARC) 2x in-plane and 2x through-plane) and 3D CUBE T2-weighted scans (sagittal; TR = 5202 ms; TE = 60 ms; flip angle = 90°; 0.8 mm isotropic voxels; matrix = 320 × 320, 226 slice; parallel imaging: ARC 2x in-plane and 2x through-plane). Functional data for resting-state and task scans were acquired with a 2D gradient-echo echo-planar imaging (EPI) sequence using simultaneous multi-slice acceleration (GE HyperBand; MB = 8): TR = 650 ms; TE = 35 ms; flip angle = 52°; 2.3 mm isotropic voxels; matrix = 92 × 92; 64 slices with 0% inter-slice gap. Spin-echo EPI field maps were also acquired with opposite phase encoding directions (Anterior-to-Posterior; Posterior-to-Anterior) and geometry matched to the functional EPI timeseries.

### MRI Preprocessing

Anatomical, functional, and resting-state data were preprocessed using *fMRIPrep*^81^ (Version 20.2.5; RRID:SCR_016216), which is based on *Nipype*^82^ (Version 1.6.1; RRID:SCR_002502), and the *ciftify* package^83^ (Version 2.3.3). The T1-weighted (T1w) image was corrected for intensity non-uniformity (INU) with *N4BiasFieldCorrection*,^84^ distributed with ANTs 2.3.3^85^ (RRID:SCR_004757), and used as the T1w-reference throughout the workflow. The T1w-reference was then skull-stripped with a *Nipype* implementation of the *antsBrainExtraction.sh* workflow (from ANTs), using OASIS30ANTs as target template. Brain tissue segmentation of cerebrospinal fluid (CSF), white-matter (WM) and gray-matter (GM) was performed on the brain-extracted T1w using *fast*^86^ (FSL 5.0.9; RRID:SCR_002823). Brain surfaces were reconstructed using *recon-all*^87^ (FreeSurfer 6.0.1; RRID:SCR_001847), and the brain mask estimated previously was refined with a custom variation of the method to reconcile ANTs-derived and FreeSurfer-derived segmentations of the cortical gray-matter of Mindboggle^88^ (RRID:SCR_002438). Freesurfer outputs were then input into *ciftify_recon_all*, which performs surface-based alignment using the MSMSulc algorithm^89^ and resamples data from native space to 32k fsLR space. The fsLR 32k space combines the benefits of nonlinear MNI volumetric registration (for subcortical structures; FSL FNIRT) with surface-level registration based on sulcal anatomy for the cortex.^90^

For each BOLD run (across all tasks and sessions), the following preprocessing was performed. First, a reference volume and its skull-stripped version were generated using a custom methodology of *fMRIPrep*. A B0-nonuniformity map (or *fieldmap*) was estimated based on two (or more) echo-planar imaging (EPI) references with opposing phase-encoding directions, with *3dQwarp*^91^ (AFNI 20160207). Based on the estimated susceptibility distortion, a corrected EPI (echo-planar imaging) reference was calculated for a more accurate co-registration with the anatomical reference. The BOLD reference was then co-registered to the T1w reference using *bbregister* (FreeSurfer) which implements boundary-based registration.^92^ Co-registration was configured with six degrees of freedom. Head-motion parameters with respect to the BOLD reference (transformation matrices, and six corresponding rotation and translation parameters) were estimated before any spatiotemporal filtering using *mcflirt*^93^ (FSL 5.0.9). The BOLD time-series were resampled onto their original, native space by applying a single, composite transform to correct for head-motion and susceptibility distortions. Gridded (volumetric) resamplings were performed using *antsApplyTransforms* (ANTs), configured with Lanczos interpolation to minimize the smoothing effects of other kernels.^94^ For the definition of TMS target regions, resting-state and pRF timeseries were analyzed in native T1 space. These native T1w timeseries were smoothed in a parcel-constrained manner (cortical ribbon and subcortical gray matter structures) with a 2mm FWHM kernel to maximize spatial accuracy of the TMS target definitions. For resting-state timeseries, this smoothing was done after additional preprocessing was performed (see *resting-state preprocessing* below).

Using the *ciftify_subject_fmri* function, pRF and spatial working memory fMRI acquisitions were then projected to the native surface in a weighted, ribbon-constrained manner, where the value assigned to each vertex is calculated as the weighted average of the voxels encompassed (or partially encompassed) by the cortical ribbon. Noisy voxels, as assessed by their local coefficient of variation, were excluded from this operation. Surface data was then resampled to fsLR 32k surface space. Subcortical data were nonlinearly resampled to MNI152 within their anatomical gray matter masks.^83,90^ Surface and volumetric data were then combined into a Connectivity Informatics Technology Initiative (CIFTI) “grayordinate” file format. Lastly, CIFTI smoothing was applied (2D surface smoothing and parcel-constrained smoothing for subcortical volumes) using a 4mm FWHM kernel. Prior to any analysis, the pRF and spatial working memory run timeseries of each voxel/vertex were detrended with a 2^nd^-order polynomial, converted to percentage signal change, and then normalized (z-score) across time points within each run.

### Resting-state preprocessing

Following fMRIPrep, we additionally performed band-pass filtering and confound regression on resting-state timeseries using the *nilearn* python package (Version 0.8.1; RRID:SCR_001362; https://doi.org/10.5281/zenodo.8397156). Several confounding time-series were calculated based on the preprocessed BOLD: framewise displacement (FD) and three region-wise global signals. FD was computed using the Power formulation^95^ (absolute sum of relative motions). The three global signals were extracted within the CSF, the WM, and the whole-brain masks. The confound time series derived from head motion estimates and global signals were expanded with the inclusion of temporal derivatives and quadratic terms for each.^96^ Frames that exceeded a threshold of 0.2 mm FD were considered motion outliers. A Butterworth temporal filter extracted frequencies between 0.009 and 0.08 Hz. The band-pass filter was also applied to all confound regressors to avoid reintroducing filtered frequency bands.^97^ The confound regression included the six head motion estimates and three region-wise global signals (CSF, WM, whole-brain), their temporal derivatives, quadratic terms, and the quadratic expansions of their derivatives (36 regressors).^96^ We additionally included spike regressors associated with motion outliers. Following confound regression, the resulting residual resting-state timeseries were *z*-scored.

### Population receptive field mapping

Stimuli were generated and presented using Python with the *PsychoPy* software package^98–100^ and were viewed with MR-compatible goggles (NordicNeuroLab VisualSystem HD). The paradigm was adapted from the procedure described in Mackey et al.^35^ Stimulus presentation was confined to a 16° × 16° field of view. The stimulus consisted of a bar aperture, which subtended 16° in length and subtended either 1°, 2°, or 3° in width. The use of different bar widths can aid the estimation of pRF size.^35^ The width of the bar aperture was held constant within each run. Functional time-series of runs consisting of the same bar width were averaged prior to pRF modeling. The bar aperture comprised 3 equally sized rectangular patches of moving dots. Dot patches swept across the visual field in a discrete manner, changing location every 1.95 s (3 TRs). The step size of each change in location was 1°. There were eight possible sweep directions: left to middle, middle to left, right to middle, middle to right, bottom to middle, middle to bottom, top to middle, and middle to top. Each sweep consisted of 6 steps/trials (11.7 s) and was followed by a 0.65 s blank fixation interval (1 TR). Each patch spanned 5.1° along the side perpendicular to the sweep direction. A 0.35° gap separated each patch. Patches of width 1°, 2°, and 3° contained 100, 200, and 300 dots, respectively. Dots had a diameter of 0.093°, moved at 1.5 °/s, and updated their position 60 times per second. At each location, observers discriminated which of the two flanking patches contained dots moving in the same direction as the middle patch. Only one of the flanker patches moved in the same direction as the middle patch on each trial. Dot motion within the middle patch was always 100% coherent. Coherent dots moved along the length of the patch opposite to the sweep direction (left or right for vertical sweeps and up or down for horizontal sweeps). The coherence of the flanker patches’ dot motion was staircased using a 1-up 3-down procedure.^101^ Moving dots had a limited lifetime of 10 frames (167 ms). Each noise dot moved in a random direction for the extent of its lifetime. Participants completed 6-9 pRF runs (either 2 or 3 repeats of each patch width).

### Spatial working memory paradigm

Spatial WM stimuli were generated and presented using Python with *PsychoPy* package.^98–100^ Each trial consisted of a sample period (325 ms), a post-sample fixation period (325 ms), a delay period with an ‘active’ or ‘drop’ cue (8.45 s), and probe period (3.25 s). Participants were instructed to maintain their gaze on a small fixation dot (0.15°) throughout the sample and delay periods. During the sample period, participants were presented with a 0.75° diameter grayscale disc at 4° eccentricity. The polar angle of the stimulus was uniformly sampled from 6 bins (0°, 60°, 120°, 180°, 240°, or 300°). ±15° of angular jitter was added randomly to the polar angle on each trial to more finely sample polar-angle space. This jitter discouraged categorical encoding of the six nominal angle bins and improved estimation of continuous spatial tuning curves, while retaining a discretized structure that ensured balanced sampling of stimulus values. As a result, 186 out of a possible 360 polar angles could be presented. Participants were not told that stimuli were limited to specific ranges and were free to report any polar angle possible (0°–359°). 325 ms following the offset of the sample stimulus, the fixation dot turned either red or blue to indicate an ‘active’ or ‘drop’ trial, respectively. On drop trials, participants did not need to maintain the item in working memory and were instructed to make a random response when probed. Following the 8.45 s delay period, subjects were presented with a probe display (3.25 s). The probe display consisted of a disc stimulus that was identical to the sample stimulus (0.75° diameter disc at 4° eccentricity). Participants used their right index, middle, and ring fingers to respond. Holding the button with their index finger moved the disc counter-clockwise along possible stimulus locations at 4° eccentricity and the ring finger moved the disc clockwise. The middle finger flipped the probe 180°. The last recorded angle was taken as the memory report for that trial. The initial angle of the probe disc was randomly generated. The randomization of the probe disc was essential to ensure that subjects were unable to prospectively plan a specific motor action (i.e., clockwise versus counter-clockwise rotation) during the delay period. As a result, we can be confident that activity over the delay-period reflects visuospatial encoding of the stimulus as opposed to motor planning. The inter-trial interval (ITI) was jittered across trials and determined using optseq2 (https://surfer.nmr.mgh.harvard.edu/optseq). 10 run sequences were generated that maximized design matrix efficiency. We specified a minimum ITI of 4.55 s (7 TRs) and a maximum ITI of 12.35 s (19 TRs) and a total run duration of 263.9 s (406 TRs). Each fMRI run comprised 14 trials with 2 repeats of each angle bin (12 active trials + 2 drop trials). Subjects completed 10 runs during the session.

In Experiment 2, participants completed a modified version of the spatial WM task on a laptop immediately following TMS. The randomization of the probe stimulus was a necessary and intentional design decision to prevent prospective motor planning. However, this aspect of the design introduces response-stage noise (e.g., the sign of behavioral error is influenced by start position and direction of rotation). Therefore, the limited trial count in Experiment 1 leads to a behavioral readout that is conservative with respect to memory precision. The time course of Experiment 2 matched the structure of Experiment 1 but was optimized for higher trial counts within the window of the cTBS effect^41^ by shortening the delay period to 1 s. Stimuli were presented at a viewing distance of approximately 60 cm. Each trial began with a 325 ms sample period in which two discs (0.75° diameter at 4° eccentricity) were shown simultaneously. One disc was red and the other was blue. After a 325 ms post-sample fixation interval, the central fixation dot changed color (blue or red) to cue which item should be retained across the delay. The probe display was identical in format to Experiment 1, and participants used the same three-button rotation response procedure to report the remembered polar angle. The initial orientation of the probe was again randomized on every trial to prevent prospective motor planning during the delay. The inter-trial interval was fixed at 0.5 s.

The task in Experiment 2 comprised 480 trials divided into 8 blocks of 60 trials, with brief self-paced breaks permitted between blocks. Trial identities were determined by a fully crossed design: six possible cued-item angle bins (0°, 60°, 120°, 180°, 240°, 300°), five angular offsets between cued and non-cued items (−120°, −60°, 60°, 120°, 180°), and two possible cued colors (red or blue). As in the fMRI experiment, ±15° of angular jitter was applied to both items on each trial, with identical jitter across the pair to preserve the specified offset.

### pRF modeling

pRF analysis was performed using the analyzePRF MATLAB toolbox.^36^ Voxel/vertex timeseries were modeled with a compressive spatial summation model,^36^ which is an extension of the pRF model described by Dumoulin & Wandell.^102^ This model includes an additional exponent parameter to account for subadditive spatial summation.^36^ The model is formally expressed as:

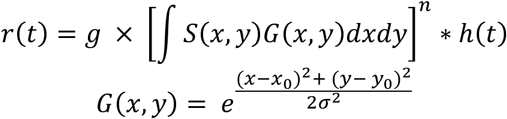

Where *r*(*t*) is a voxel’s predicted response at timepoint *t*, *g* is a gain parameter, *S* is a binary stimulus mask, *G* is a 2-dimensional isotropic Gaussian, *n* is an exponent parameter controlling the degree of subadditivity (≤1), *h*(*t*) is a canonical hemodynamic response function, *x*_0_ and *y*_0_ are parameters defining the position of the Gaussian, and *σ* is a parameter defining the standard deviation of the Gaussian. Prior to fitting, functional time-series of runs consisting of the same experimental bar size (1, 2, or 3°) were averaged. To train the pRF model to predict responses to the spatial working memory task, we further averaged the time-series of each bar width across subjects.^37,103^ The original pRF model described by Dumoulin and Wandell^102^ involved a two-stage fitting procedure: an initial coarse grid-fit followed by an exhaustive non-linear optimization procedure using seed parameters from the grid-fit. However, recent studies have demonstrated that the full optimization does not outperform the grid-fit when predicting independent, left-out data.^35^ It was argued that the grid-fit procedure is more robust to noise and better able to predict the responses of fronto-parietal voxels with large pRFs. We have previously used the grid-fit procedure to identify retinotopically organized cerebellar areas.^22^ The grid-fitting procedure iterated over 49,045 possible parameter combinations (25 eccentricities, 24 angles, 17 widths, and 5 exponents) to find the grid point that produced the maximal correlation between the predicted and actual voxel timeseries. Predicted and actual timeseries were polynomial detrended and scaled to unit length prior to computing this correlation. The gain or amplitude parameter was estimated following the grid-fit procedure as:

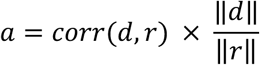

where *d* is the voxel timeseries, *r* is the predicted timeseries associated with the maximal correlation grid point, and ∥ ∥ denotes vector length. Note that there is an interaction between the width of the Gaussian and the static nonlinearity parameter such that pRF size effectively equals 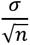 as:

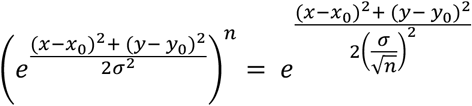

### TMS target definition

We first identified a cerebellar lobule VIIb target in each hemisphere. To do so, we projected the lobule VIIb retinotopic mask from the van Es cerebellar retinotopy atlas^37^ into the subject’s native T1 space. We then identified the voxel within this mask with the maximal pRF R^2^ value, a pRF eccentricity <8°, and a polar angle preference ±45° from the ipsilateral hemifield horizontal meridian. If no voxel could be identified with these criteria, we removed the eccentricity criterion. If this also failed to identify a voxel within the lobule VIIb mask, we further removed the polar angle range criterion. Following the identification of a lobule VIIb target in each hemisphere, we created an 8 mm sphere centered on this same voxel and then applied a cerebellar cortical ribbon mask. This ROI was used as a seed in seed-based functional connectivity analyses to identify IPS, sPCS, and somatosensory control TMS targets. Due to the crossing of cortico-cerebellar circuits, left hemisphere cerebellar lobule VIIb was used as a seed for right hemisphere cerebral cortex targets, and right hemisphere cerebellar lobule VIIb was used as a seed for left hemisphere cerebral cortex targets. We defined anatomical search spaces for each cortical target. For the IPS target, we used the IPS0 label from the Wang probabilistic retinotopy atlas.^104^ For the sPCS target, we combined the FEF and area 6a labels from the Glasser atlas.^90^ Lastly, for the somatosensory control site, we combined the postcentral gyrus and paracentral gyrus/sulcus labels from the Destrieux atlas,^105^ and then manually trimmed this label to lie medial to the hand knob portion of postcentral gyrus and posterior to the central sulcus. For each subject, these labels were projected from the *fsaverage* template surface to that subject’s native surface (*mri_label2label*). The native surface label was then projected into native T1 volume space in a cortical ribbon constrained manner (*mri_label2vol*). IPS and sPCS targets were defined as the voxels with the maximal correlation with contralateral retinotopic cerebellar lobule VIIb. The somatosensory control target was identified as the voxel with the *minimum* absolute correlation with the contralateral cerebellar lobule VIIb retinotopic ROI.

### TMS protocol

Repetitive TMS (rTMS) was delivered through a MagPro X100 magnetic stimulator and a 70 mm figure-8 coil (MC-B70, MagVenture). In an initial screen session, motor evoked potentials (MEPs) elicited using biphasic posterior-anterior stimulation and coil oriented 45° to the coronal plane were recorded from the left first dorsal interosseous (FDI) using surface electromyography (2-channel built-in EMG device, Rogue Research, Montreal). Active motor threshold (AMT) was recorded as the percentage of stimulator output that elicited an MEP of ≥ 200 μV peak-to-peak on five out of ten trials while the participant was contracting the FDI muscle at 20% of maximum.^106,107^ At the end of the screening session, participants were given several pulses at 80% AMT over regions of the scalp roughly corresponding to cerebellar lobule VIIb, IPS, SPCS, and the control site to ensure that the stimulation intensity would be tolerable in future sessions. One participant reported discomfort related to stimulation of the approximate sPCS site and as a result only received lobule VIIb, IPS, and control stimulation in subsequent sessions. In the experimental sessions following the baseline fMRI session, we delivered continuous theta-burst stimulation (cTBS) with standard parameters: 3 pulses of stimulation at 50 Hz, repeated every 200 ms, for a total of 600 pulses over 40 seconds. We used the Brainsight Frameless neuronavigation system (Rogue Research, Montreal CA) to align the previously acquired structural scans to each participant’s skull in real-time to allow for precise targeting.

TMS thresholds have been shown to depend on the distance between the coil and underlying target stimulation site.^108^ To account for between-site differences in scalp-to-cortex distance, we adjusted stimulation intensity for each target relative to primary motor cortex (M1). Specifically, we (1) measured scalp-to-cortex distance at the M1 hand knob and at each target site (lobule VIIb, IPS, sPCS, and control) from each participant’s T1-weighted anatomical image, and (2) computed a distance-adjusted active motor threshold as

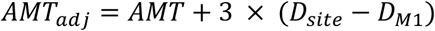

where *D*_site_ and *D*_M1_ denote the scalp-to-cortex distance (cm) at the target site and the M1 hand knob, respectively. Continuous theta-burst stimulation was delivered at 80% of *AMT*_adj_ for each site.

### Regions of Interest

ROIs were defined based on the baseline pRF mapping and atlas definitions. The anatomical search spaces for visual and parietal ROIs were extracted from a probabilistic retinotopy atlas (V1-V3, V3AB, IPS0-3).^104^ We used the ‘FEF’ label from the HCP MMP atlas^90^ for our sPCS ROI. A cerebellar retinotopic atlas was used to define the anatomical search space for lobule VIIb.^37^ For visual cortex ROIs, we selected vertices with a pRF variance explained greater than 20% and an eccentricity between 3° and 8°. For the remaining ROIs in frontoparietal cortex and cerebellum, which possess larger receptive field sizes, we selected vertices with a pRF variance explained greater than 20% and no eccentricity criterion.

### Spatial encoding model

To characterize the effect of TMS applied to each target on spatial selectivity throughout the brain, we employed a generative model-based, probabilistic decoding approach. This method was initially developed to characterize neural uncertainty associated with the encoding of orientation in visual cortex.^38^ This method has been extended to model the encoding of motion direction^23^ and spatial polar angle.^40^ The model specifies a generative distribution, which represents the probability that a particular stimulus *s* will evoke an activation pattern *b* (i.e., *p*(*b*|*s*)). This distribution takes the form of a multivariate normal distribution with a stimulus-dependent mean vector *f*(*s*) and a voxel covariance matrix Ω. The mean vector *f*(*s*) is determined by each voxel’s tuning profile over possible working memory stimulus locations (0°-359° polar angle at 4° eccentricity). Here, rather than use a linear channel-based approach as in prior work,^23,38–40,109^ we instead use the nonlinear pRF model trained on the mapping data collected during the baseline fMRI session as our encoding model. The *i*-th voxel’s spatial tuning curve was modeled as:

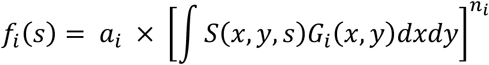

where *a_i_* is the *i*-th voxel’s gain estimated from the independent pRF mapping session, *S*(*x*, *y*, *s*) is a binary stimulus mask of stimulus *s* (0.75° disc at 4° eccentricity, 0°–359° polar angle) spanning the entire 16° × 16° display, *G_i_*(*x*, *y*) is a 2D Gaussian representing *i*-th voxel’s pRF, and *n_i_* is the *i*-th voxel’s estimated static nonlinearity parameter.

The regularized estimation of the noise covariance had two components: shrinkage of the sample correlation matrix towards a simpler structure comprising noise correlations related to tuning similarity and global noise shared across all voxels, as well as shrinkage of the sample variance toward its median value.

The noise correlation shrinkage target was constructed as follows. A sample covariance matrix was first computed from residuals resulting from the comparison of model-predicted responses (*f*(*s*)) and the training set, and converted to a correlation matrix:

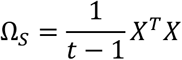

where *X* is a *t* × *v* matrix of residuals (where *t* is the number of trials in the training set and *v* is the number of voxel/vertices). The sample correlation matrix was then computed as:

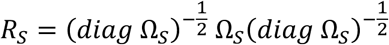

We then computed voxel tuning covariance as:

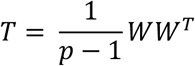

Where *p* is the number of display pixels, and *W* represents unwrapped voxel pRFs (2D Gaussians) for each voxel forming a *v* × *p* matrix. This covariance matrix was converted to a correlation matrix:

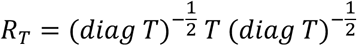

The global noise correlation was computed by regressing the empirical sample correlations onto tuning similarity and an intercept. The intercept captures the noise common to all voxels. The noise correlation shrinkage target was then computed as:

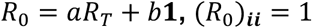

Where *a* is the slope parameter relating the tuning correlation matrix with the sample correlation matrix, *b* is the intercept, and **1** is a *v* × *v* matrix of ones. (*R*_0_)***_ii_*** = 1 ensures that the resulting matrix is a valid correlation matrix (ones along the diagonal). Sample correlations were then shrunk toward the target correlation matrix, with the degree of shrinkage controlled by *λ*.

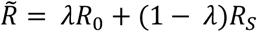

We additionally shrank voxel variance toward the median variance across voxels:

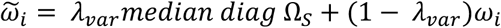

where *λ_var_* is the parameter controlling the degree of shrinkage toward the median and *ω_i_* is the sample variance of the *i*-th voxel. Thus, the noise covariance around voxel spatial tuning curves can be computed as:

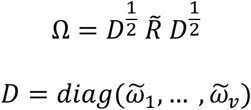

Thus, the generative distribution of voxel/vertex responses follows a multivariate distribution with a mean *f*(*s*) and covariance Ω:

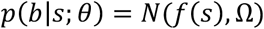

where *θ* = {*x*_0_, *y*_0_, *σ*, *a*, *n*, *λ*, *λ_var_*} are the model’s parameters.

Model fitting and assessment was performed using a 10-fold leave-one-run-out cross-validation scheme, which iteratively partitioned the data into a training set and a test set. Trial-wise delay-period responses were estimated by averaging the final three TRs of the delay period, corresponding to the last 1.95 s before probe onset. This late-delay window was chosen to maximize separation from the sample-evoked response while avoiding contamination from probe- and response-related activity. Because the probe had not yet appeared during this window, these responses could not reflect processing of the probe stimulus or execution of the report. Prior to cross-validation, runs were reordered to align stimulus sequences across participants and then averaged trial-wise across participants to create a “super-subject.” This approach follows prior large-scale pRF work^37,103^ in which fMRI time series are aligned across subjects, averaged, and then modeled to recover stable group-level parameters. This procedure also has the advantage of allowing stimulation-order effects to be averaged across counterbalanced participants before estimating condition-specific spatial tuning. We assumed pRF centers (*x*_0*i*_, *y*_0*i*_) were fixed across tasks (spatial-attention motion-discrimination pRF task and spatial working memory task) and TMS targets (CB, IPS, SPCS, control), but allowed voxel-wise modulation of baseline (additive), gain (multiplicative), and width (multiplicative) by task/target. Within each outer training fold, we performed a grid-based posterior approximation per voxel with log-uniform gain ∈ [2^−3^, 2^3^] in 0.25-log2 steps, log-uniform width ∈ [2^−2^, 2^2^] in 0.25-log2 steps, and baseline ∈ [−0.20,0.20] in 0.01 steps (17,425 combinations). Assuming flat priors over grid points, we converted log-densities to probabilities, normalized, and took the posterior mean of each parameter to regenerate *f*(*s*) for 0° − 359°.

We then performed an inner 9-fold cross-validation to estimate noise covariance hyperparameters (*λ*, *λ_var_*). This procedure comprised a coarse grid search (*λ* ∈ (0,1] and *λ_var_* ∈ [0, 1]) followed by a fine grid search around the point associated with the minimum loss.^110^ At each grid point, we evaluated the negative log-likelihood (scaled by 2/v) of the validation sample covariance (*S_val_*) under a zero-mean Gaussian model with covariance Ω:^111^

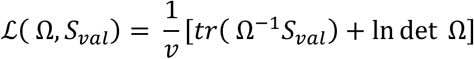

where constant terms independent of Ω have been dropped.^111^ Hyperparameters associated with the minimum cross-validated loss were then used to re-estimate Ω for the entire training set (9 runs).

Then, for each trial in the test set, we used Bayes rule to obtain a posterior probability distribution over spatial polar angle (0°–359°), indicating which spatial location was most probable given the observed pattern of BOLD responses.

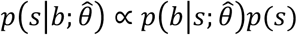

Where *θ̂* are the estimated model parameters. The stimulus prior, *p*(*s*), was set to 1 for all polar angles as experimental stimuli were uniformly distributed. The posterior mean represented the decoded location and the posterior standard deviation represented the decoded uncertainty. Using the posterior mean, we calculated mean circular squared error (MSE_circ_). The inverse of MSE_circ_ produced a measure of estimation efficiency, with higher values associated with more accurate estimation of spatial location.^112^ To test for differences in decoded uncertainty, we computed the mean posterior standard deviation across trials for each ROI and stimulation condition.

### Tuning curve modulation

To characterize the effects of stimulation on voxel response profiles, we performed a “tuning-curve modulation” analysis that estimates per-voxel modulation of gain (multiplicative), width (multiplicative scaling of pRF size), baseline response (additive), and response variance (multiplicative) across TMS conditions. This analysis uses the same encoding model and notation introduced above for spatial decoding, but is conducted outside the cross-validation procedure. The center coordinates (*x*_0*i*_, *y*_0*i*_) of each voxel’s pRF and the power-law nonlinearity parameter (*n_i_*) were held fixed across tasks and TMS sessions.

We modeled the session-dependent modulation of voxel tuning curves with a parameter vector

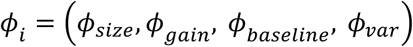

where *ϕ_size_* multiplicatively scales the pRF width, *ϕ*_*gain*_ multiplicatively scales gain, *ϕ_baseline_* additively scales the baseline, and *ϕ_var_* multiplicatively scales the voxel’s response variance. For each voxel *i*, we generated a library of candidate tuning curves by scaling the pRF width in 2D display space and then projecting to polar angle. Specifically, we replaced σ_i_ with *ϕ_size_* σ_i_ inside the Gaussian *G_i_*(*x*, *y*) defined over the 16° × 16° display, evaluated the 2D integral with the stimulus mask *S*(*x*, *y*, *s*) to obtain 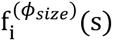. In contrast, the remaining modulation parameters operated on the resulting 1D tuning curve across polar angle. The modulated tuning curve used in the likelihood was

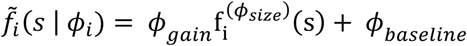

with stimulus-independent variance *ϕ_var_v̂_i_* (where *v̂_i_* is the *i^th^* voxel’s variance estimate from the baseline pRF-mapping residuals).

Let *r_ti_* denote the measured delay-period response of voxel *i* on working memory trial *t*, and *s_t_* represent the presented polar angle stimulus on that trial. Assuming conditionally independent Gaussian noise,

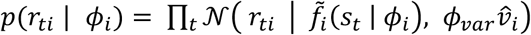

We evaluated the log-likelihood on a dense Cartesian grid with uniform priors:

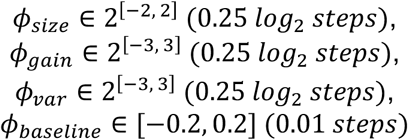

Posterior weights were obtained by exponentiating log-likelihoods after subtracting the maximum log-likelihood to avoid numerical underflow, and then normalizing to sum to one. With uniform priors on the grid, posterior means for each parameter were computed as weighted averages under these normalized weights. We then regenerated each voxel’s modulated tuning curve across polar-angle,

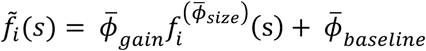

with *ϕ̅_var_v̂_i_* as the response variance.

### Statistical Analysis

All analyses were implemented in R (Version 4.5.1). All comparisons used non-parametric permutation tests with two-sided *p*-values from 5,000 permutations. Unless noted, contrasts were made relative to the somatosensory control site. False-discovery rate (Benjamini–Hochberg) corrections were applied once per family: 3 comparisons for behavioral analyses (CB, IPS, or sPCS versus Control) and 30 comparisons for fMRI analyses (10 ROIs × 3 session contrasts). Permutation *p*-values were computed as (#{∣ *T*_perm_ ∣≥∣ *T*_obs_ ∣} + 1)/(*N*_perm_ + 1)^113^ and then FDR-adjusted.

Secondary comparisons among the WM stimulation sites were conducted using a gated follow-up procedure. For each fMRI measure, the primary family of 30 active-versus-control contrasts was tested and FDR-corrected as described above. If at least one WM active stimulation condition differed significantly from control within a given ROI, all three pairwise comparisons among the active sites (CB versus IPS, CB versus sPCS, and IPS versus sPCS) were then performed for that ROI. FDR correction was applied across the complete set of active-site comparisons performed for that measure. These follow-up tests used the same permutation unit and test statistic as the corresponding active-versus-control analysis, with the pairwise activesite difference substituted as the contrast of interest.

To test for differences in spatial working memory recall error across TMS conditions, trials were first screened to remove likely guesses as on random-guess trials the error distribution is approximately uniform, so error magnitude ceases to index precision (e.g., 45° versus 90° error both reflect failed encoding for a guess rather than a graded loss of precision). In Experiment 1, trials with absolute error greater than the mean absolute error plus two standard deviations (computed within subject and session) were excluded (Control: 4.17% ± 0.73% of trials excluded; CB: 4.44% ± 0.85%; IPS: 2.62 ± 0.32%; sPCS: 5.28% ± 0.88%). In Experiment 2, the same error criterion was applied and trials with reaction time > mean RT plus two standard deviations (reaction time was not recorded in Experiment 1) were additionally excluded (Control: 7.15% ± 0.57% of trials excluded; CB: 6.74% ± 0.56%; IPS: 6.36% ± 0.61%; sPCS: 6.81% ± 0.36%). Session effects were then tested with a hierarchical paired (within-subject) permutation procedure in which, for each contrast, session labels were randomly swapped within subject, subject-level means recomputed, and the across-subject mean paired difference used as the test statistic. To account for potential sequence effects, we additionally regressed out session order using a robust linear mixed-effects model with a random intercept per subject (*robustlmm* package),^114^ and repeated the permutation test on the residualized errors.

Differences in decoding metrics were assessed separately for estimation efficiency and decoded uncertainty. For estimation efficiency, the predicted polar angle was permuted across trials before recomputing the inverse mean circular squared error within each ROI and session. Decoded uncertainty, quantified as the mean posterior standard deviation, was tested by permuting session labels across trials within each ROI before recomputing session means.

Primary contrasts compared each active stimulation session with the control session. Where the gated follow-up criterion was met, the same procedures were used to test pairwise differences among the active stimulation sessions.

Session effects on tuning-curve modulation were tested separately for each posterior-mean modulation parameter: gain, baseline, pRF width, and residual variance. Session labels were permuted across voxels within each ROI, and the mean difference between each active stimulation condition and control stimulation was recomputed. Where the gated follow-up criterion was met for a given parameter and ROI, the same voxel-level permutation procedure was used for pairwise comparisons among the active stimulation sites.

## Data Availability

Summary behavioral and fMRI datasets underlying the statistical analyses reported in this study will be publicly available through the Open Science Framework (OSF) at https://osf.io/2b6um. Owing to the large size of the raw MRI datasets and associated storage constraints, raw neuroimaging data are not included in the public repository but may be made available by the corresponding author upon reasonable request.

## Code Availability

Code used for modeling, statistical analysis, and figure generation will be made publicly available at https://github.com/brissend/CB_WM_TMS.

## Acknowledgements

We thank T. Polk, J. Jonides, T. Adkins, Q. Nguyen, J. Sellers, and R. Panda for helpful comments and discussion. This work was supported by NIH grant no. F32MH124268 (J.A.B.). The funder had no role in the study design, data collection and analysis, decision to publish or preparation of the manuscript.

## Author Contributions Statement

J.A.B. and T.G.L. conceived the study. J.A.B. collected the data. J.A.B. carried out data analysis and wrote the original draft of the manuscript. All authors reviewed the manuscript and provided critical revisions. T.G.L. and M.V. provided resources and supervision.

## Competing Interests Statement

The authors declare no competing interests.

---

**Supplementary Table 1.** Probabilistic Spatial Decoding Results.

| ROI | Contrast | Estimation Efficiency |  | Posterior SD |  |
| --- | --- | --- | --- | --- | --- |
| | | $\Delta$ (deg<sup>-2</sup>) | $p_{FDR}$ | $\Delta$ (°) | $p_{FDR}$ |
| V1 | Control – CB | 1.83e-05 | 0.24 | -2.84 | 0.43 |
|  | Control – IPS | **-4.10e-05** | **0.01** | 2.43 | 0.47 |
|  | Control – sPCS | -2.27e-05 | 0.17 | 4.30 | 0.19 |
| V2 | Control – CB | 2.86e-05 | 0.06 | -3.88 | 0.17 |
|  | Control – IPS | -1.01e-05 | 0.50 | 3.12 | 0.21 |
|  | Control – sPCS | 1.40e-05 | 0.34 | 3.09 | 0.18 |
| V3 | Control – CB | -6.00e-06 | 0.70 | 0.96 | 0.78 |
|  | Control – IPS | -5.21e-06 | 0.70 | **6.32** | **0.03** |
|  | Control – sPCS | **3.89e-05\*** | **0.01** | 0.49 | 0.86 |
| V3AB | Control – CB | **3.34e-05\*** | **0.02** | **-9.65\*** | **0.0004** |
|  | Control – IPS | 2.61e-05 | 0.10 | **-9.37\*** | **0.0004** |
|  | Control – sPCS | 2.02e-05 | 0.25 | **-10.55\*** | **0.0004** |
| IPS0 | Control – CB | **3.63e-05\*** | **0.01** | -4.26 | 0.13 |
|  | Control – IPS | 1.88e-05 | 0.25 | **-11.52\*** | **0.0004** |
|  | Control – sPCS | 8.99e-06 | 0.52 | -0.82 | 0.78 |
| IPS1 | Control – CB | -8.33e-07 | 0.94 | **-24.11\*** | **0.0004** |
|  | Control – IPS | 1.59e-05 | 0.27 | -5.15 | 0.13 |
|  | Control – sPCS | 1.67e-05 | 0.25 | **-24.70\*** | **0.0004** |
| IPS2 | Control – CB | -1.26e-05 | 0.40 | **16.86** | **0.0004** |
|  | Control – IPS | -1.70e-05 | 0.25 | **9.19** | **0.02** |
|  | Control – sPCS | 4.75e-06 | 0.71 | **18.11** | **0.0004** |
| IPS3 | Control – CB | -9.65e-06 | 0.50 | **-9.00\*** | **0.03** |
|  | Control – IPS | -1.67e-05 | 0.25 | **-14.35\*** | **0.0004** |
|  | Control – sPCS | 5.33e-06 | 0.70 | **-19.07\*** | **0.0004** |
| sPCS | Control – CB | 1.66e-05 | 0.25 | **9.51** | **0.002** |
|  | Control – IPS | 1.86e-05 | 0.25 | **34.45** | **0.0004** |
|  | Control – sPCS | 1.23e-05 | 0.40 | **5.53** | **0.03** |
| VIIb | Control – CB | 1.10e-05 | 0.4463318 | **-40.17\*** | **0.0005** |
|  | Control – IPS | 3.38e-06 | 0.7737974 | **-42.85\*** | **0.0005** |
|  | Control – sPCS | 5.60e-06 | 0.6909133 | **-18.10\*** | **0.0005** |

\* – Deficit relative to control stimulation; Bolded text – $p < 0.05$ corrected

**Supplementary Table 2.**
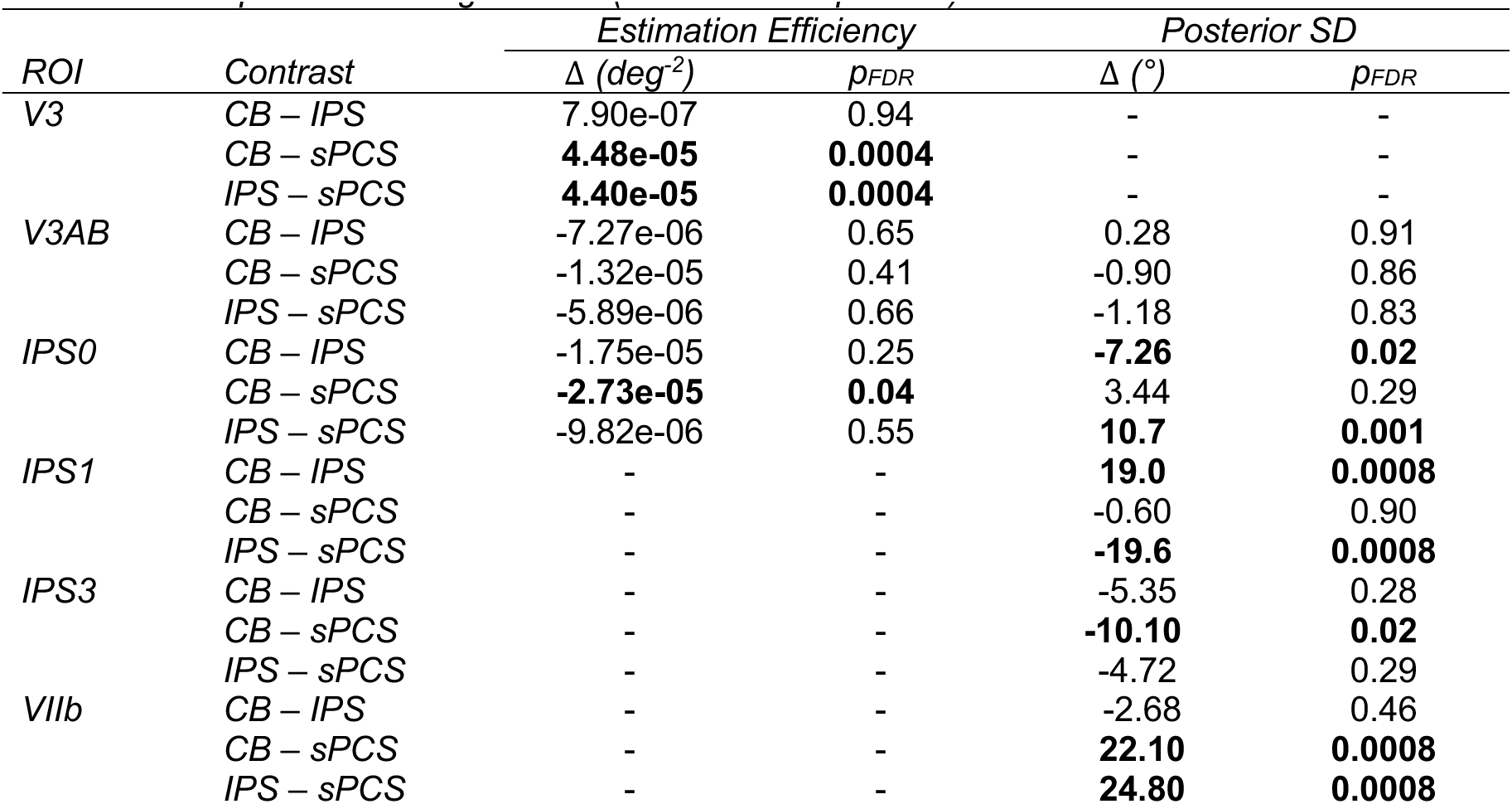
Probabilistic Spatial Decoding Results (WM Site Comparison)

**Supplementary Table 3.** Spatial Tuning Modulation Results.

| ROI | Contrast | $\phi_{\text{baseline}}$ | | $\phi_{\text{gain}}$ | | $\phi_{\text{size}}$ | | $\phi_{\text{var}}$ | |
| --- | --- | --- | --- | --- | --- | --- | --- | --- | --- |
| | | $\Delta$ | $p_{\text{FDR}}$ | $\Delta$ | $p_{\text{FDR}}$ | $\Delta$ | $p_{\text{FDR}}$ | $\Delta$ | $p_{\text{FDR}}$ |
| V1 | Control – CB | **0.02\*** | **0.0002** | **0.24\*** | **0.0004** | 0.02 | 0.46 | **0.03** | **0.0003** |
|  | Control – IPS | **0.05\*** | **0.0002** | 0.05 | 0.23 | **0.21** | **0.0008** | -0.006 | 0.220 |
|  | Control – sPCS | **0.02\*** | **0.0002** | **-0.22** | **0.001** | 0.01 | 0.67 | **0.03** | **0.0003** |
| V2 | Control – CB | **0.01\*** | **0.0004** | **0.21\*** | **0.002** | **0.06** | **0.01** | **-0.006\*** | **0.01** |
|  | Control – IPS | **0.04\*** | **0.0002** | **-0.19** | **0.01** | **0.13** | **0.0008** | **-0.02\*** | **0.0003** |
|  | Control – sPCS | **0.02\*** | **0.0002** | -0.12 | 0.20 | 0.04 | 0.09 | **-0.01\*** | **0.0003** |
| V3 | Control – CB | **0.01\*** | **0.0002** | **0.25\*** | **0.0008** | 0.02 | 0.41 | **0.01** | **0.04** |
|  | Control – IPS | **0.04\*** | **0.0002** | -0.08 | 0.413 | -0.01 | 0.81 | **-0.04\*** | **0.0003** |
|  | Control – sPCS | **0.03\*** | **0.0002** | -0.12 | 0.198 | 0.04 | 0.08 | -0.006 | 0.08 |
| V3AB | Control – CB | **0.02\*** | **0.0002** | **0.32\*** | **0.0004** | -0.05 | 0.116 | **0.03** | **0.0003** |
|  | Control – IPS | **0.03\*** | **0.0002** | **0.32\*** | **0.0004** | -0.04 | 0.109 | **-0.05\*** | **0.0003** |
|  | Control – sPCS | **0.03\*** | **0.0002** | **0.16\*** | **0.003** | 0.03 | 0.408 | **-0.102\*** | **0.0003** |
| IPS0 | Control – CB | **0.02\*** | **0.0002** | **0.11\*** | **0.003** | 0.03 | 0.304 | **-0.04\*** | **0.005** |
|  | Control – IPS | **0.02\*** | **0.0002** | **0.21\*** | **0.0004** | **-0.08\*** | **0.0007** | -0.01 | 0.541 |
|  | Control – sPCS | **0.03\*** | **0.0002** | -0.04 | 0.35 | **0.06** | **0.0388** | **-0.07\*** | **0.0003** |
| IPS1 | Control – CB | **0.02\*** | **0.0002** | **0.29\*** | **0.0005** | -0.01 | 0.78 | **-0.105\*** | **0.0003** |
|  | Control – IPS | **0.02\*** | **0.0002** | **0.09\*** | **0.04** | **0.07** | **0.0008** | **0.0860** | **0.0003** |
|  | Control – sPCS | **0.03\*** | **0.0002** | **0.27\*** | **0.0004** | 0.02 | 0.45 | **0.102** | **0.0003** |
| IPS2 | Control – CB | **0.04\*** | **0.0002** | **0.12\*** | **0.0004** | **0.151** | **0.0008** | **0.03** | **0.02** |
|  | Control – IPS | **0.03\*** | **0.0002** | **0.13\*** | **0.0004** | 0.0251 | 0.180 | **0.09** | **0.0003** |
|  | Control – sPCS | **0.03\*** | **0.0002** | 0.02 | 0.45 | **0.168** | **0.0008** | **0.04** | **0.002** |
| IPS3 | Control – CB | **0.03\*** | **0.0002** | **0.24\*** | **0.0004** | **-0.08\*** | **0.03** | **-0.09\*** | **0.0003** |
|  | Control – IPS | **0.03\*** | **0.0002** | **0.22\*** | **0.0004** | 0.02 | 0.65 | -0.001 | 0.97 |
|  | Control – sPCS | **0.02\*** | **0.0002** | **0.23\*** | **0.0004** | **-0.07\*** | **0.02** | **-0.05\*** | **0.002** |
| sPCS | Control – CB | **0.002\*** | **0.003** | -0.005 | 0.61 | 0.006 | 0.554 | **-0.125\*** | **0.0003** |
|  | Control – IPS | **0.004\*** | **0.0002** | -0.03 | 0.08 | **0.05** | **0.003** | **0.133** | **0.0003** |
|  | Control – sPCS | -0.001 | 0.09 | -0.008 | 0.44 | **-0.02\*** | **0.02** | **-0.289\*** | **0.0003** |
| VIIb | Control – CB | **0.01\*** | **0.0002** | **0.32\*** | **0.0004** | **-0.14\*** | **0.0008** | **-0.07\*** | **0.009** |
|  | Control – IPS | **0.03\*** | **0.0002** | **0.46\*** | **0.0004** | **-0.13\*** | **0.0008** | **-0.21\*** | **0.0003** |
|  | Control – sPCS | **0.01\*** | **0.0002** | **0.257\*** | **0.0004** | **-0.08\*** | **0.03** | -0.001 | 0.97 |

\* – Deficit relative to control stimulation; Bolded text – $p < 0.05$ corrected

**Supplementary Table 4.**
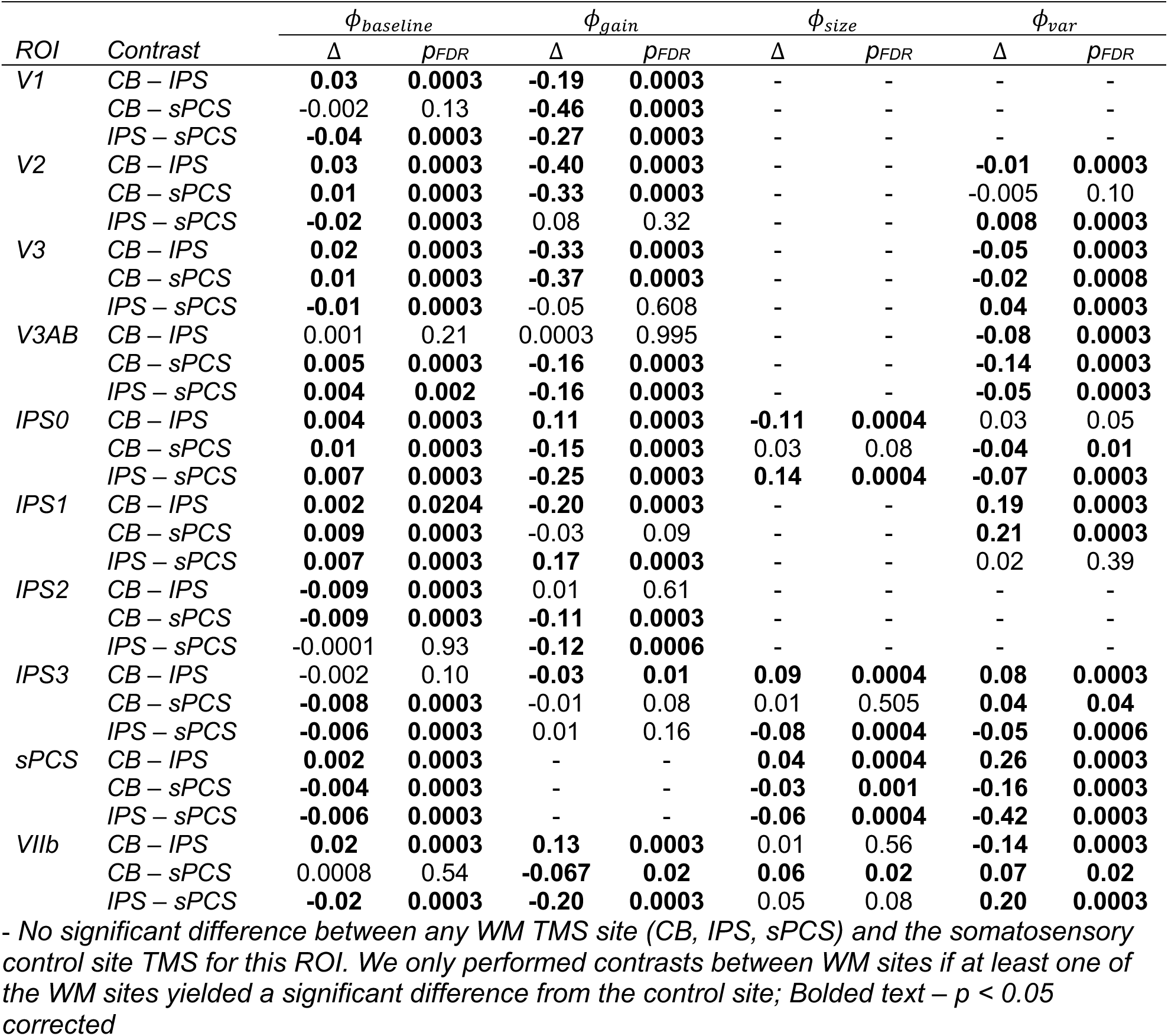
Spatial Tuning Modulation Results (Working Memory Site Comparison)

| ROI | Contrast | $\phi_{baseline}$ | | $\phi_{gain}$ | | $\phi_{size}$ | | $\phi_{var}$ | |
| --- | --- | --- | --- | --- | --- | --- | --- | --- | --- |
| | | $\Delta$ | $p_{FDR}$ | $\Delta$ | $p_{FDR}$ | $\Delta$ | $p_{FDR}$ | $\Delta$ | $p_{FDR}$ |
| V1 | CB – IPS | **0.03** | **0.0003** | **-0.19** | **0.0003** | - | - | - | - |
|  | CB – sPCS | -0.002 | 0.13 | **-0.46** | **0.0003** | - | - | - | - |
|  | IPS – sPCS | **-0.04** | **0.0003** | **-0.27** | **0.0003** | - | - | - | - |
| V2 | CB – IPS | **0.03** | **0.0003** | **-0.40** | **0.0003** | - | - | **-0.01** | **0.0003** |
|  | CB – sPCS | **0.01** | **0.0003** | **-0.33** | **0.0003** | - | - | -0.005 | 0.10 |
|  | IPS – sPCS | **-0.02** | **0.0003** | 0.08 | 0.32 | - | - | **0.008** | **0.0003** |
| V3 | CB – IPS | **0.02** | **0.0003** | **-0.33** | **0.0003** | - | - | **-0.05** | **0.0003** |
|  | CB – sPCS | **0.01** | **0.0003** | **-0.37** | **0.0003** | - | - | **-0.02** | **0.0008** |
|  | IPS – sPCS | **-0.01** | **0.0003** | -0.05 | 0.608 | - | - | **0.04** | **0.0003** |
| V3AB | CB – IPS | 0.001 | 0.21 | 0.0003 | 0.995 | - | - | **-0.08** | **0.0003** |
|  | CB – sPCS | **0.005** | **0.0003** | **-0.16** | **0.0003** | - | - | **-0.14** | **0.0003** |
|  | IPS – sPCS | **0.004** | **0.002** | **-0.16** | **0.0003** | - | - | **-0.05** | **0.0003** |
| IPS0 | CB – IPS | **0.004** | **0.0003** | **0.11** | **0.0003** | **-0.11** | **0.0004** | 0.03 | 0.05 |
|  | CB – sPCS | **0.01** | **0.0003** | **-0.15** | **0.0003** | 0.03 | 0.08 | **-0.04** | **0.01** |
|  | IPS – sPCS | **0.007** | **0.0003** | **-0.25** | **0.0003** | **0.14** | **0.0004** | **-0.07** | **0.0003** |
| IPS1 | CB – IPS | **0.002** | **0.0204** | **-0.20** | **0.0003** | - | - | **0.19** | **0.0003** |
|  | CB – sPCS | **0.009** | **0.0003** | -0.03 | 0.09 | - | - | **0.21** | **0.0003** |
|  | IPS – sPCS | **0.007** | **0.0003** | **0.17** | **0.0003** | - | - | 0.02 | 0.39 |
| IPS2 | CB – IPS | **-0.009** | **0.0003** | 0.01 | 0.61 | - | - | - | - |
|  | CB – sPCS | **-0.009** | **0.0003** | **-0.11** | **0.0003** | - | - | - | - |
|  | IPS – sPCS | -0.0001 | 0.93 | **-0.12** | **0.0006** | - | - | - | - |
| IPS3 | CB – IPS | -0.002 | 0.10 | **-0.03** | **0.01** | **0.09** | **0.0004** | **0.08** | **0.0003** |
|  | CB – sPCS | **-0.008** | **0.0003** | -0.01 | 0.08 | 0.01 | 0.505 | **0.04** | **0.04** |
|  | IPS – sPCS | **-0.006** | **0.0003** | 0.01 | 0.16 | **-0.08** | **0.0004** | **-0.05** | **0.0006** |
| sPCS | CB – IPS | **0.002** | **0.0003** | - | - | **0.04** | **0.0004** | **0.26** | **0.0003** |
|  | CB – sPCS | **-0.004** | **0.0003** | - | - | **-0.03** | **0.001** | **-0.16** | **0.0003** |
|  | IPS – sPCS | **-0.006** | **0.0003** | - | - | **-0.06** | **0.0004** | **-0.42** | **0.0003** |
| VIIb | CB – IPS | **0.02** | **0.0003** | **0.13** | **0.0003** | 0.01 | 0.56 | **-0.14** | **0.0003** |
|  | CB – sPCS | 0.0008 | 0.54 | **-0.067** | **0.02** | **0.06** | **0.02** | **0.07** | **0.02** |
|  | IPS – sPCS | **-0.02** | **0.0003** | **-0.20** | **0.0003** | 0.05 | 0.08 | **0.20** | **0.0003** |

- No significant difference between any WM TMS site (CB, IPS, sPCS) and the somatosensory control site TMS for this ROI. We only performed contrasts between WM sites if at least one of the WM sites yielded a significant difference from the control site; Bolded text – $p < 0.05$ corrected

**Supplementary Figure 1.**
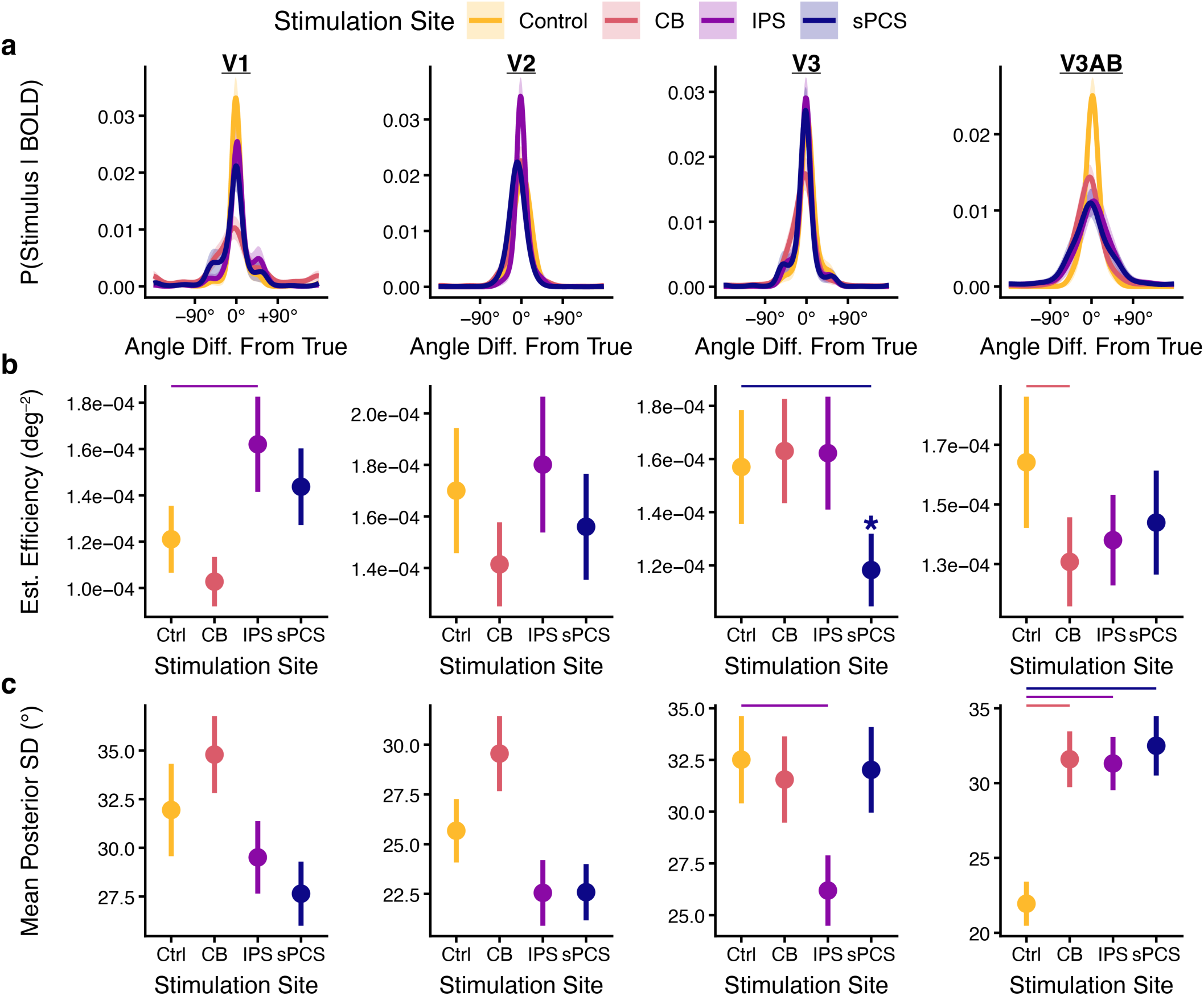
Visual cortex spatial decoding. (a) Mean posterior probability distribution. Averaging trial-aligned posterior distributions produces a session-level probability density over stimulus locations for each region of interest (ROI). Sharper peaks around 0° reflect better decoding of the remembered spatial location. (b) Estimation efficiency (inverse of mean circular squared error between the posterior mean and the true stimulus) reflecting decoding accuracy for each session and region. (c) Mean posterior standard deviation reflecting decoding uncertainty. Error bars reflect a bootstrap estimate of standard error. Each column reflects results for one ROI.

**Supplementary Figure 2.**
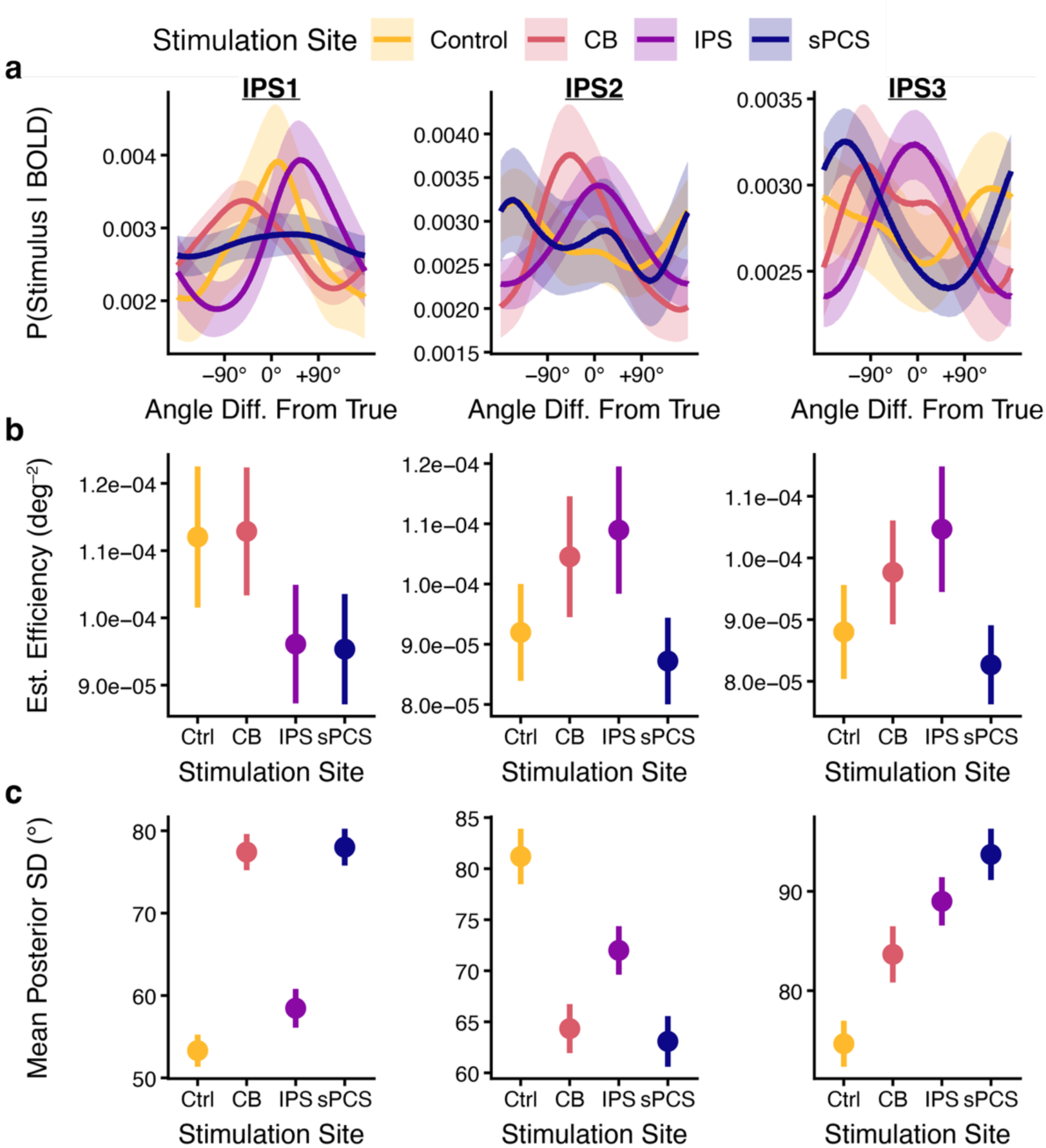
Parietal cortex spatial decoding. (a) Mean posterior probability distribution. Averaging trial-aligned posterior distributions produces a session-level probability density over stimulus locations for each region of interest (ROI). Sharper peaks around 0° reflect better decoding of the remembered spatial location. (b) Estimation efficiency (inverse of mean circular squared error between the posterior mean and the true stimulus) reflecting decoding accuracy for each session and region. (c) Mean posterior standard deviation reflecting decoding uncertainty. Error bars reflect a bootstrap estimate of standard error. Each column reflects results for one ROI.

## Notes

### Competing Interest Statement

The authors have declared no competing interest.

### Summary of Updates

Minor revisions to text and figures

